# An active torque dipole across tissue layers drives avian left-right symmetry breaking

**DOI:** 10.1101/2025.07.16.665037

**Authors:** Julia Pfanzelter, Jonas Neipel, Adrian A. Lahola-Chomiak, Nikoloz Tsikolia, Alexander Mietke, Jerome Gros, Frank Jülicher, Stephan W. Grill

## Abstract

Unlike in mice, frogs, and fish, left-right (L/R) body axis formation in avian embryos does not arise from the chiral beat of cilia. Instead, a counter-clockwise tissue rotation around Hensen’s node, the organizer of amniote development, repositions cells expressing L/R sidedness genes. Yet, the physical origin of this rotation remains elusive. Here, we provide evidence that in quail embryos, the node tissue generates an active chiral torque of ∼6µNµm to drive the chiral tissue flow. Microsurgery experiments reveal that this torque depends on actomyosin molecular activity, is generated within the dorsal node tissue, and requires the underlying ventral meso-/endoderm to act as a mechanical substrate sustaining the counter-torque. We conclude that a dorsoventrally oriented tissue-scale active torque dipole at the node translates cell-scale chirality to organismal L/R asymmetry, adding a mechanical dimension to the canonical function of embryonic organizers as signaling hubs.

## Introduction

From subatomic particles to whole organisms, physical systems across scales exhibit intrinsic handedness, or chirality [1, 2]. During embryonic development, the microscopic mechanical activity of motor molecules acting on chiral cytoskeletal filaments, i.e. actin or microtubules, translates molecular chirality to an organismal scale [3–5]. In mouse, fish and frog embryos, microtubule-based cilia drive a directional extracellular fluid flow initiating a signaling cascade to establish left–right (L/R) asymmetry and body plan handedness [6–10]. However, in many vertebrates—including birds, reptiles, and some mammals—this cilia-based mechanism is absent [11–16].

In avian embryos, L/R asymmetry instead arises from a chiral tissue flow: the epiblast rotates counterclockwise around Hensen’s node, repositioning areas of gene expression to break L/R symmetry and pattern the L/R body axis [11, 12, 17–21]. Mechanical activity of the actomyosin cortex is required for this process [11, 22], but how cellular forces coordinate to produce the chiral, tissue-scale rotation remains unknown.

The spontaneous generation of chiral flows from microscopic mechanical activity is a hallmark of active chiral matter [23–27]. In *C. elegans*, the active chiral nature of the actomyosin cortex lies at the heart of L/R body axis formation: active torque dipoles generated within the cortex at molecular scales drive chiral flows of cell surfaces, which, at the 4-6 cell stage of the embryo, facilitate the generation of a L/R asymmetric cell-cell contact pattern [25, 28–30]. Also in other invertebrates, actomyosin components such as actin, myosin and formin have been implicated in L/R symmetry breaking [31–38]. Furthermore, *in vitro* studies have revealed, that vertebrate cells also exhibit an actomyosin-dependent chirality [39–43]. However, it is not well understood if cell-scale chiral mechanical processes play a role in L/R body axis formation of vertebrate embryos consisting of thousands of cells. Specifically, it is unclear if the chiral tissue rotation is the consequence of cell-scale active torque generation, or if it results from already existing L/R asymmetries at the tissue-scale without the need of a chiral mechanical process. This distinction is central to understanding how microscopic chirality propagates across scales to contribute to tissue-level dynamics during vertebrate development.

## Results

### Quantification of chiral tissue flow

It has been previously reported that the Hensen’s node in chick embryos (*Gallus gallus*) undergoes a collective rotation at the time the primitive streak starts to regress towards the posterior [11, 12]. We first set out to determine if the node in quails undergoes a similar rotation. We performed *ex ovo* live imaging of 25 transgenic quail embryos (*Coturnix coturnix japonica*) ubiquitously expressing a fluorescent membrane marker, and quantified tissue movement of the epiblast with particle image velocimetry (PIV) from 10 hours prior to 10 hours post streak regression onset. We find that while the epiblast undergoes large-scale flows towards the streak at this time, [18, 44–49] (Fig. 1d,e, Movie 1), it also undergoes a characteristic rotation in a counter-clockwise direction similar to chick (Fig. 1e). In addition, as in chick [17, 18], the anterior tip of the streak develops a prominent leftward kink (Fig. 1g,k, S3). We next quantified the rotation rate of the central node tissue together with a net leftward velocity, both obtained with respect to the surrounding epiblast tissue (Eq. S7,S8). Neither the rotation rate nor the leftward velocity is significantly different from zero at early times (Fig. 1i,j; Δ*t <* −5 h; see also Fig. S1 and S4). Between 5 hours prior and 3 hours post streak regression onset (*t* = 0h), the node performs a counter-clockwise rotation when viewed from the dorsal side, and the node rotation rate peaks with 5.4 (4.9, 6.1) deg */h* at *t* = 0h (Fig. 1i), where here and in the following *X* (*Y, Z*) denotes the median *X* and its confidence interval in terms of the (5^th^, 95^th^) percentile from bootstrapping (see SI 1.2.9). In addition, the node moves towards the left between 2 hours prior and 5 hours post streak regression onset (Fig. 1j). Together, these tissue flows result in a leftward kinking of the anterior tip of the streak by *θ*_kink_ = 37° (32°, 42°) (Fig. 1k). To understand the spatial extent of chiral tissue rotation, we considered the flow field in a reference frame of the epiblast tissue surrounding the node and decomposed the flow field into its mirror-symmetric and mirror-antisymmetric components (Fig. 1c,l,m,n). This reveals significant chiral tissue flow up to a distance of 150*µ*m away from the node center (Fig. 1m,n, S4). We conclude that the tissue in the vicinity of the node undergoes a collective counter-clockwise rotation and a leftward displacement at the time of L/R symmetry breaking.

**Fig. 1.**
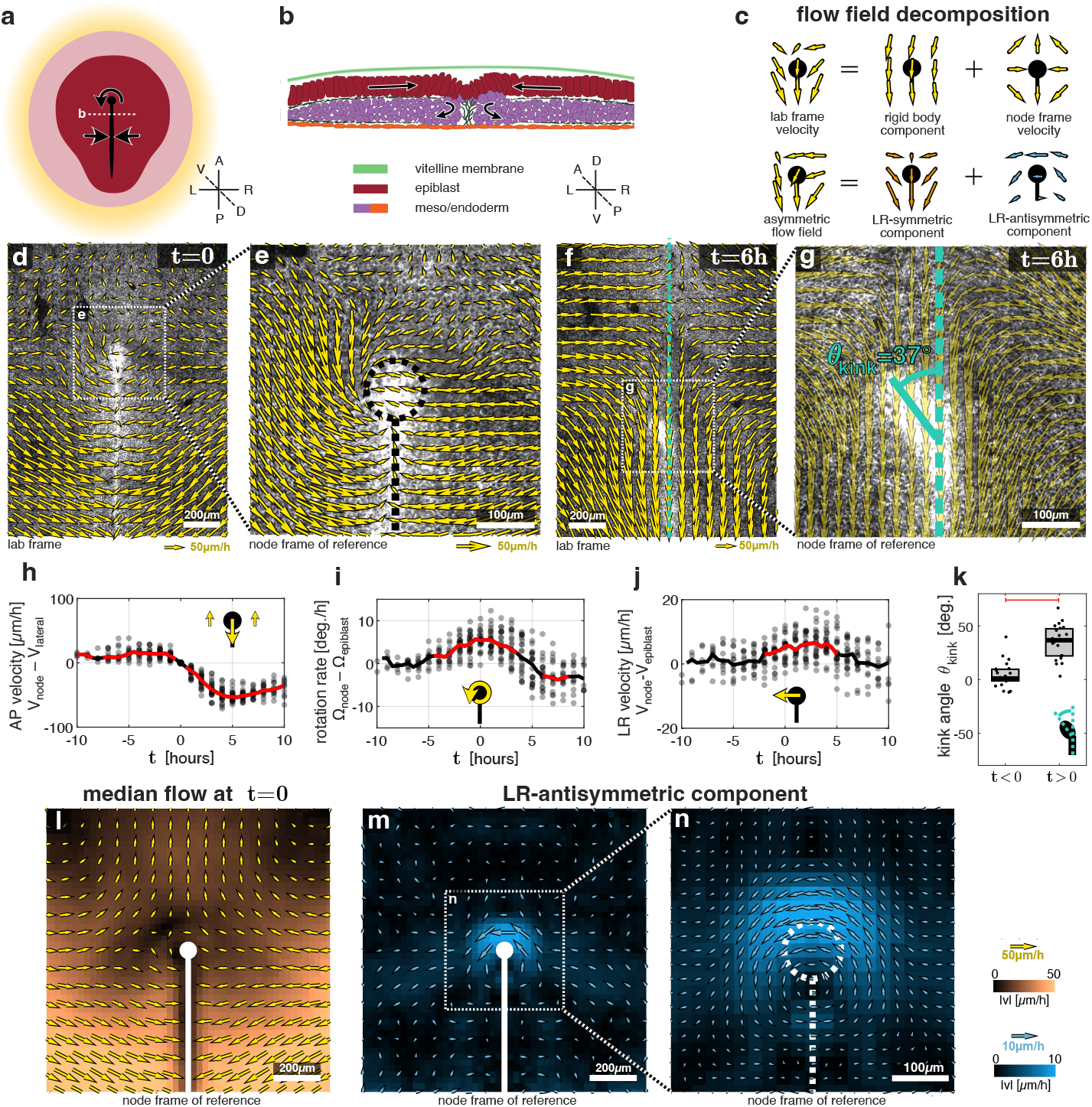
Quantification of chiral tissue flow. **a,b:** Schematic of a quail embryo at the time point of L/R symmetry breaking (stage HH4+, maximum streak extension) with tissue flows indicated by black arrows. **a:** dorsal view. **b:** cross-section of the primitive streak (black line in **a**) along the white dashed line in **a. c:** Schematic of flow field decompositions, which we used to quantify chiral tissue flow. Except in **d**,**f**, we always consider the node frame velocity, i.e. the flow field in the reference frame of the tissue surrounding the node (see SI 1.2.4 for details).s **d-g:** microscopy images of the epiblast of a developing quail embryo expressing a fluorescent membrane marker overlaid with the flow field obtained by PIV (same embryo as in Movie 1). **e** and **g** are a zoom-in of the white dashed box in **d** and **f** respectively. Black dashed line in **e** indicates the streak and node region. Turquoise dashed line in **f**,**g** denotes the embryo mid-line relative to which the kink angle *θ*_kink_ is quantified as shown in **g**. *t* denotes the timing with respect to the onset of streak regression (*t* = 0) (see Fig. S2). **h-j:** Quantification of tissue flow in terms of the translation and rotation of the node relative to the surrounding tissue (see SI 1.2.3 and 1.2.5). Black dots indicate 1h time windows in single embryos. Solid line indicates sample median with red color denoting time points where the median is significantly different from 0 (*p <* 0.01, Wilcoxon signed rank test). **k:** box plot of kink angle of the node as in **g**, using the maximal angle for time points before (*t <* 0) and after (*t >* 0) the onset of streak regression (see SI 1.2.7 and Fig. S3). Red line indicates significant difference with *p <* 0.01 (Wilcoxon rank sum test). **l:** Embryo-median of tissue flow around the node (white circle) at *t* = 0 (see Fig. S4 for other time points). **m**,**n** L/R antisymmetric component of median tissue flow, where **n** is a zoom-in of the white dashed box in **m**.

### Inferring active forces from measured tissue flow

We next set out to shed light onto the mechanical basis of this collective chiral tissue flow. We describe the epiblast tissue as a 2D active fluid in which flows are driven by active force generation at the molecular scale through e.g. the activity of molecular motors [50, 51]. We next inferred the active force density driving symmetric and antisymmetric tissue flows by determining a second-order spatial derivative of the respective flow fields (Fig. 2a-d and see SI, Eq S19). We find that, on the one hand, symmetric tissue flow is associated with active forces localized all along the streak (Fig. 2a,b and Fig. S6). These amount to a contraction of the streak tissue along the L/R axis, and drive the large-scale tissue flow towards the streak, where cells leave the epithelium as part of gastrulation ([44, 49, 52–54], Fig. 1b). On the other hand, antisymmetric tissue flow is associated with active forces localized only to a region within 100µm of the node center, peaking anterior to the node center (Fig. 2c,d). When integrated over this region, these active forces amount to a net torque of ∼ 6*µ*N*µ*m exerted with respect to the node center (SI, Eq. S38; using an effective 2D tissue shear viscosity of *η* = 1*mNh/m* [55]). This indicates that the node tissue drives its own rotation by generating a net torque, which must be generated with respect to a substrate, i.e. the underlying meso/endoderm or the vitelline membrane above. Such a coupling to a mechanical substrate implies a finite hydrodynamic length 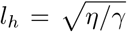 where *γ* is an effective friction coefficient arising from the interactions with a substrate [56]. We infer *l*_*h*_ from a boundary analysis (see SI 2.2, Fig. S6,S7). When analyzing L/R-symmetric flows, we find a large hydrodynamic length *l*_*h*_ *>* 300*µ*m (Fig. S6), consistent with previous studies [44, 52, 53]. Remarkably, when analyzing L/R-antisymmetric flows, we find a significantly reduced hydrodynamic length *l*_*h*_ ∼ 50 − 100*µ*m, (Fig. S7). This is consistent with a picture where mechanical coupling to an underlying substrate is relevant for chiral flows, while it is not for the large scale flows towards the streak (see SI 2.6). A possibility is that the chiral tissue flow is driven by a tissue-scale dorsoventrally oriented active torque dipole localized at the node, rotating the epiblast tissue against the underlying meso-/endoderm. In contrast, the large scale flows towards the streak result from force dipoles within the epiblast, effectively driving tissue flow of both layers [57]. In conclusion, our results suggest that the node tissue drives its own rotation by generating a net torque with respect to an underlying substrate.

**Fig. 2.**
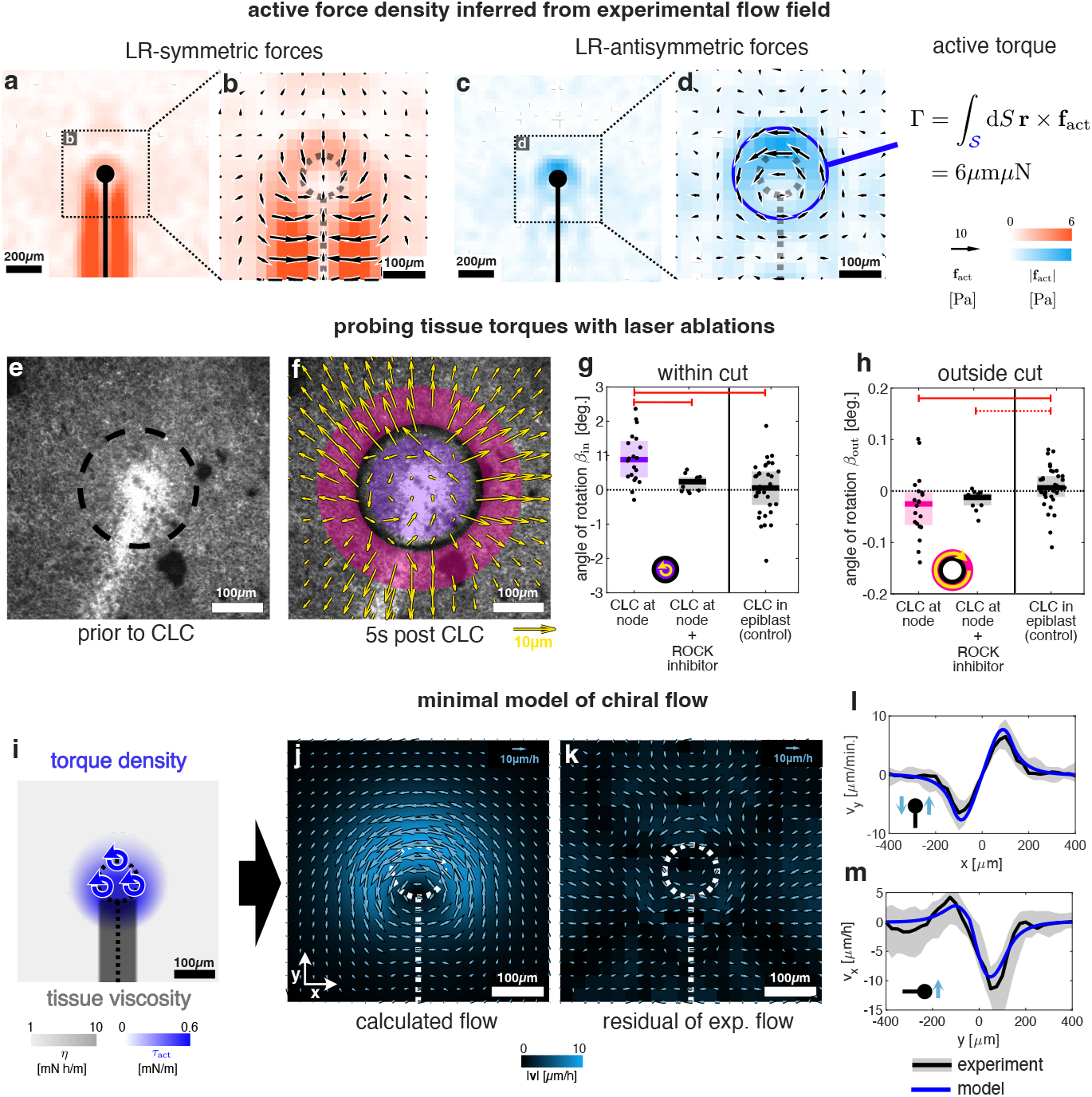
A torque at the node drives the chiral tissue flow. **a-d:** active force densities **f**_act_ (Eq. S19) indicated by black arrows with color of the background indicating the magnitude. Forces in **a**,**b** underlying large-scale L/R-symmetric tissue flows were calculated from the median L/R symmetric flow (Fig. S4) at *t* = 0 using *η* = 1mN h*/*m, *l*_*h*_ *→ ∞* [55]. Forces in **c**,**d** underlying chiral tissue flow were calculated from the median L/R antisymmetric flow (Fig. 1m) using *η* = 1mN h*/*m, *l*_*h*_ = 75*µ*m (see [55] and SI 2.2.4 and 2.3). The blue circle in **d** with radius 100*µ*m indicates the integration area for obtaining the active torque Γ that the node tissue generates.**b** and **d** are a zoom-in of the dashed box in **a** and **c** respectively **e**,**f** microscopy images of the quail epiblast before (**e**) and after (**f**) circular laser ablation (CLC) as shown in Movie 2. Black dashed line indicates where the tissue was cut. Yellow arrows indicate the displacement field, purple area and pink area indicate the analyzed cut area and surrounding area respectively. **g** and **h**: angles of rotation calculated from the displacement field in the cut area and the surrounding area, respectively, in the presence and absence of the ROCK inhibitor (H1152). Positive angles correspond to a counter-clockwise rotation, i.e. in the direction of chiral flow, such that *β*_in_ *>* 0 reveals a torque generated at the node. Note also that we do not observe a net tissue displacement upon laser ablation (Fig. S11). Red solid (dashed) lines indicate statistical significance with *p <* 0.01 (*p <* 0.05) comparing conditions using a two-sided Wilcoxon rank sum test **i:** spatial fields underlying the active fluid model of chiral tissue flow inferred from the measured flow field (see SI 2.5). Blue shading denotes the amplitude of the active torque density *τ*_act_ modeled in terms of the first two harmonic modes on a circular domain (Eq. S42,S44), whereas gray shading denotes the magnitude of the tissue viscosity *η*. **j**: flow field of an active chiral fluid (**i**) as a model for chiral tissue flow, plotted analogous to Fig. 1n. **k**: residual of the experimental flow (Fig. 1n) after subtracting the model flow in **j. l:** Plot of the spatial profile of the AP tissue velocity (*v*_*y*_) as function of the L/R position relative to the midline (*x* = 0) across a line through the node comparing the model flow field (blue as in **j**) with the experimentally measured flow field (black line indicating the embryo median as in Fig. 1m with gray area indicating the 25th to 75th percentile). **m:** Plot of the spatial profile of the L/R velocity11(*v*_*x*_) across a line through the streak (*y <* 0) and the node (*y* = 0) analogous to **l**.

### Laser ablation reveals actomyosin-dependent torque at the node

We next set out to provide direct evidence for local torque generation at the node. In the past, stresses in living matter have been revealed by quantifying shear deformations on the elastic time scale upon laser ablation [56, 58–62]. Here, we probe active torque generation by performing circular laser cuts (CLC). If the node generates an active torque, then abruptly isolating the node by CLC should lead to an immediate rotation of the node tissue in the direction of chiral tissue flow (SI 2.4). If torque generation is limited to the node, then the surrounding tissue should rotate in the opposite direction following CLC. We performed CLC around the node and quantified the immediate angle of rotation *β*_in_ of the tissue inside the circular cut region, together with the immediate angle of rotation *β*_out_ in a ring surrounding the cut region, over the subsequent 5 s using PIV (see SI 1.2.8 and Movie2). We find that for CLC centered around the node *β*_in_ = 0.88° (0.48°, 1.30°) and *β*_out_ = −0.03° (−0.06°, −0.01°), *n* = 20, while there are on average no significant rotations for CLC in the epiblast away from the node (*β*_in_ = 0.05° (−0.18°, 0.41°) and *β*_out_ = 0.01° (0.00°, 0.02°); *n* = 34, Fig. 2e-h). The rapid counterclockwise rotation of the node and the clockwise rotation of the surrounding epiblast tissue following laser ablation reveal a tissue-scale torque rotating the node counterclockwise. We conclude that a tissue-scale dorsoventrally oriented active torque dipole drives the rotation of the dorsal node tissue. This rotation is slowed down by the surrounding epiblast tissue.

The cortical actomyosin cytoskeleton of cells has been demonstrated to generate active torques [25, 30, 43, 63]. We next asked if actomyosin is responsible for active torque generation at the node [11]. We used the ROCK inhibitor H1152 to inhibit actomyosin mechanical activity, and pursued CLC at the node as before to probe active torque generation. We find that the node rotation *β*_in_ upon CLC is significantly reduced when applying 25 µM ROCK inhibitor for 1.5-2 hours (Fig. 2g,h, *β*_in_ = 0.24° (−0.04°, 0.37°) and *β*_out_ = −0.01° (−0.03°, 0.00°), *n* = 11). We conclude that torque generation at the node depends on actomyosin mechanical activity.

### An active fluid model of chiral tissue flow

We hypothesize that dorsoventrally oriented active torque dipoles at the cellular scale together generate the tissue-scale active torque dipole at the node [23–25, 29, 63, 64]. We thus wanted to determine the distribution of active torque dipoles at the node that can account for our measured L/R antisymmetric active force density (Fig. 2d) and tissue-scale chiral flow (Fig. 1n). We consider the streak tissue to be more rigid than the surrounding epiblast in order to capture the experimentally observed net leftward tissue flow (Fig. 2i, see SI 2.5). With this, we find that an AP-asymmetric torque dipole density field localized to the node region drives a tissue flow field that is in agreement with the experiment up to a residual of the order of embryo to embryo variation (Fig. 2j-m; SI 2.5). The associated torque density is on the order of 10^−16^Nm*/*(*µm*)^2^, consistent with chiral torques measured in single cells [63]. Integrating the calculated tissue flow field over a time interval of 6 hours yields an angle of rotation *θ*_flow_ of the node that is on the order of the corresponding experimental measurements (*θ*_flow_ = 39° as compared to *θ*_flow_ = 30°(26°, 32°), see SI, Eq. S11). In addition, placing a line along the streak and advecting it with the calculated flow field over the same time period yields a kinking angle of *θ*_kink_ = 45° (Fig. S9), consistent with a picture where the leftward kinking of the anterior tip of the streak results from chiral tissue flow (Fig. 1g,k). We conclude that a L/R symmetric distribution of dorsoventrally oriented active torque dipoles at the node is sufficient to account for the observed chiral tissue flow and kinking of the streak.

### Mechanical coupling to ventral tissue facilitates chiral tissue flow

Our results up to now indicate that the node tissue generates a torque to drive its own rotation. Angular momentum conservation implies that the node tissue must be mechanically coupled to a substrate to sustain the counter-torque. We hypothesize that the underlying ventral meso-/endoderm tissue acts as said substrate (Fig. S10). Using an eye-brow knife, we mechanically removed the entire ventral tissue 2-5 hours before the onset of streak regression, to evaluate if both chiral tissue flow and leftward kinking are subsequently reduced. While embryos underwent streak regression (16 out of 25, see SI table S1) at normal speeds following ventral tissue removal (Fig. S5), both node rotation and leftward kinking were significantly reduced (*θ*_flow_ = 21° (15°, 28°) and *θ*_kink_ = 12° (7°, 19°); Fig. 3d,g,h, movie 3). Note that the ventral tissue reforms over time such that 5 hours after the perturbation, the meso-/endoderm tissue beneath the node is approximately as thick as in unperturbed embryos (Fig.3j-l). Consistent with this observation, node rotation is not entirely abolished upon removal of the ventral tissue, but significantly reduced and delayed (Fig. 3h,i). In addition, replacing the ventral tissue with a transplanted vitelline membrane rescues tissue rotation and kink formation (*θ*_flow_ = 31° (27°, 36°) and *θ*_kink_ = 29° (21°, 39°) (Fig.3e,g,h, movie 4). This demonstrates that a passive ventral substrate can sustain a counter torque in a manner that is sufficient for node rotation. Taken together, our results are consistent with a picture where a direct mechanical coupling between epiblast and ventral tissue enables active torque generation, driving chiral tissue flow and node rotation (Fig. 3m).

**Fig. 3.**
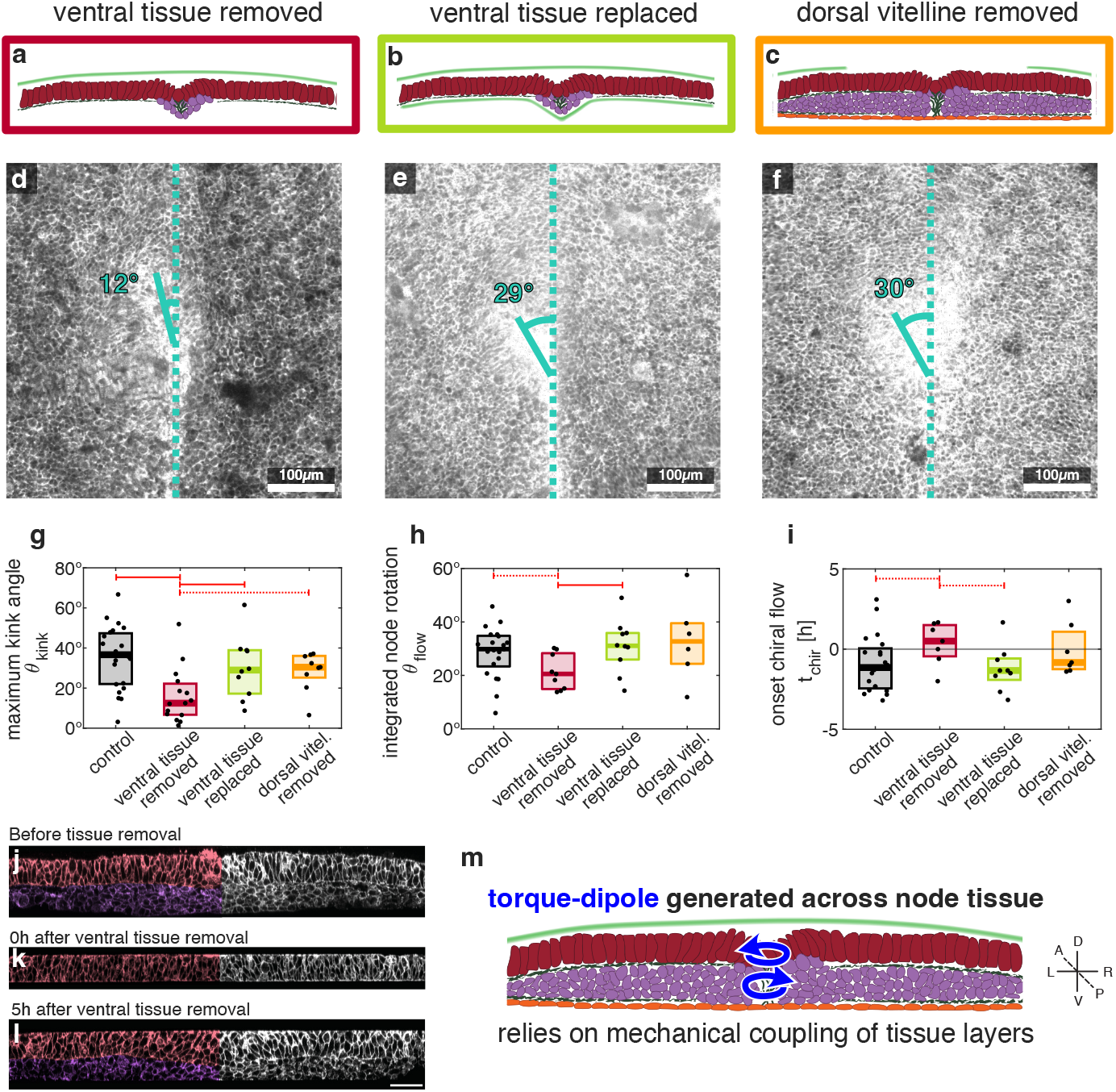
Chiral flow relies on mechanical coupling to ventral tissue. **a-c:** sketches of quail embryos analogous to Fig. 1b after microsurgery where ventral tissue layers were removed **a** or replaced by a vitelline membrane **b**, or where the dorsal vitelline membrane was removed around the node **c. d-f:** microscopy images of quail epiblast (snapshots of Movie 3 to Movie 5) at *t* = 6*h* analogous to Fig. 1g following the perturbations sketched in **a-c**, respectively. Kink angle indicated in turquoise as in Fig. 1g. Numbers are sample medians as indicated in **g. g-i:** box plots of leftward kinking (**g** as in Fig. 1k for *t >* 0), integrated node rotation (**h**, see Eq S11 quantifying instantaneous rotation as in Fig. 1i) and onset of chiral flow (**i**, onset defined as half maximum of node rotation (Ω_node_ Ω_epiblast_), timing with respect to onset of streak regression at *t* = 0). Perturbation indicated by color of box as in **a**-**c** with statistics of unperturbed embryos shown in gray. Black dots indicate single embryos. Red solid (dashed) lines indicate statistical significance with *p <* 0.01 (*p <* 0.05) comparing conditions using a one-sided Wilcoxon rank sum test, which shows that ventral tissue removal perturbs and delays chiral flow and leftward kinking. Angles upon removal of vitelline membrane: *θ*_flow_ = 33° (24°, 47°), *θ*_kink_ = 30° (27°, 36°). Median values for onset of chiral flow: control: *t*_chir_ = −1.2*h* (−2.3*h*, −0.2*h*); ventral tissue removed: *t*_chir_ = +0.5*h* (−0.6*h*, +2.6*h*); ventral tissue replaced: *t*_chir_ = −1.3*h* (−1.7*h*, −0.7*h*); dorsal vitel. removed: *t*_chir_ = −0.8*h* (−1.3*h*, +1.5*h*). **j-l** LR-DV cross-sections of fixed embryos centered at the streak with left half of epiblast colored in red and meso-/endoderm in violet analogous to Fig. 1b. In **j**, a control embryo is shown whereas in **k** and **l** the ventral tissue was removed just before fixation **k** or 5h prior to fixation **l**, where we observe recovery of the ventral (meso-/endoderm) tissue. Scale bar = 50*µ*m. **m**: illustration of the mechanics underlying chiral flow inferred from the results in Fig. 2 and **g-i**. Blue arrows indicates torques. Note that these rotational forces are not equivalent to rotational flows. For example, if the ventral tissue violet) is effectively rigid, the dorso-ventrally oriented torque dipole drives rotational flows only in the dorsal epiblast (red).

## Discussion

The L/R body axis in avian embryos is established via a collective tissue rotation [11, 12], which we here show to result from an actomyosin-dependent chiral active mechanical process: an active torque dipole across Hensen’s node drives a chiral tissue rotation against the underlying meso-/endoderm substrate.

It will be interesting to investigate the 3D nature of chiral flows. On one hand it will be informative to determine if the torque dipole also results in a counter-rotation of the meso-/endoderm, or if the meso-/endoderm is effectively rigid. On the other hand, resolving the structure of the Hensen’s node at the (sub)cellular level can reveal how a supracellular active torque dipole between tissue layers results from cell-scale active chiral processes. Such an understanding of the translation of chirality across scales is relevant far beyond the avian Hensen’s node: Multiple studies have shown that cells across the animal kingdom exhibit an actomyosin-dependent chirality [25, 30, 31, 33, 35–43], but so far it remains unclear under which circumstances such a cell-scale chirality translates to tissue-scale chiral morphogenesis and the establishment of a L/R body axis. Furthermore, we note that many vertebrate species that do not appear to rely on cilia for L/R symmetry breaking have a primitive streak–like structure [11, 13– 16, 65]. It is therefore tempting to speculate that the generation of an active torque dipole across tissue layers is a general mechanism for L/R body axis establishment, relevant to various amniote species.

## Acknowledgments

We thank Aurelien Villedieu for experimental advice and helpful discussions; Teije Middelkoop, Patrick Breier and Arjun Narayanan for helping start this project; Marko Popovic and Francis Corson for insightful discussions, especially regarding the the-oretical aspects. We thank Katrin Reppe and the Animal Facility as well as Britta Schroth-Diez and the Light Microscopy Facility at MPI-CBG for their support; and Anne Grapin-Botton and Carl Modes for comments on the manuscript.

