## Supplemental Information for "An active torque dipole across tissue layers drives avian left-right symmetry breaking"

July 15, 2025

**3 Supplementary Figures** **24**

**4 Supplementary movies** **32**

### 1 Methods

#### 1.1 Experimental Methods

##### 1.1.1 Embryo Culture

The transgenic quail line expressing a GFP membrane marker [Tg(hUbC:memGFP)] was a generous gift from Dr. Jérôme Gros [67]. Quail embryos were cultured using the modified EC culture system [66]. Quail eggs were gathered from the on-site facility by biomedical services and stored at 14°C. Eggs were incubated at 38°C for 2-3 hours before dissection. Cleaned embryos were then cultured on growth media (50% loose albumen, 0.16% Glucose, 0.1M NaCl, 0.2% Agarose [Roth 2267.4]) overnight at 38°C until stage HH4. All protocols were conducted in consultation with the animal welfare officer.

##### 1.1.2 Live Microscopy

For live imaging the Olympus IX83 Spinning Disc Confocal Microscope equipped with a Yokogawa CSU-W1 scan head was used. Embryos mounted on a thin layer of growth media in a glass-bottomed 6-well plate (CellVis P06-1.5H-N) were incubated at 38°C and ambient air. Samples were illuminated with a 488nm laser using an Olympus U Plan XApo 20x (0.8 NA) air objective and signal detected with a Hamamatsu ORCA-Fusion BT Digital CMOS camera (SN 500674/0). The system is operated via Olympus cellSens software version 4.1.

##### 1.1.3 Chemical treatment

For the drug delivery experiment 25µM of the ROCK inhibitor H1152 dihydrochloride (Tocris) were added to the growth medium and embryos were cultured for 1.5 - 2 hours in the presence of the inhibitor prior to laser ablation experiments.

##### 1.1.4 Laser ablation

Circular laser cuts were performed on a spinning-disk confocal microscope (Zeiss Axio Observer.Z1, 10x Apo 0.45NA, Yokogawa CSU-X1 scan head, Zeiss AxioCam 705 Mono, CCD camera) using the 355 nm laser (1 kHz rep rate, max 42 $\mu$J pulse power, 1 ns pulse length) from RAPP Optoelectronic to cut circles with a radius between (50 - 150$\mu$m). Due to software limitation defining the direction of cut (clockwise vs counterclockwise) consisted of a clockwise cut immediately followed by a counterclockwise cut through the already cut region or vise-versa. The sample was heated with a stage-top incubator and the system was operated with ZEN 3.2 Blue and SysCon2.

##### 1.1.5 Fixed Microscopy

To visualize tissue architecture, embryos were fixed in 4% PFA dissolved in 1x PBS for 30min then mounted under a coverslip immersed in VectashieldPlus (Vector laboratories Ref H-1900-10) using scotch tape as spacer. A Leica DMI 4000, inverted stand microscope with a Leica 63x/1.3 HC PLAPO CS 2, Glycerine, objective was used. Samples were illuminated with a 955nm tuneable laser and signal detected with a Leica spectral detection HyD detector.

##### 1.1.6 Scraping Microdissection and Rescue

To remove the ventral cell layers embryos were cultured as describe above until HH4. An eyebrow knife was prepared by mounting a human eyebrow hair onto a glass Pasteur pipette using melted paraffin wax. Drops of HBSS were applied to the ventral side of the embryo and the eyebrow was used to gently scrape away the meso-/endoderm. Excess fluid was then removed and embryos were cultured as normal. To rescue this phenotype a segment of vitelline membrane from another egg located far away from the embryo was attached to a filter paper ring. This piece was then overlayed onto an embryo, where the ventral meso-/endoderm was removed such that the inner leaf of the vitelline membrane was in contact with the ventral side of the embryo.

##### 1.1.7 Cutting a hole in the vitelline membrane

Embryos were placed dorsal side up onto a metal ring, 20-50 µl of FastGreen were placed onto the vitelline membrane and using a sharpened sewing needle a hole was created in the vitelline membrane on top of the Hensen's node. Afterwards, excess liquid was removed and embryos were placed dorsal side down on the agar pad and cultured as usual.

#### 1.2 Analysis of experimental data

##### 1.2.1 Data selection

Across conditions, we successfully performed live-imaging of 89 embryos. However, some embryos exhibited aberrant phenotypes leading to the premature death of the embryo. In particular we observed ruptures of the streak in several embryos. We excluded embryos where such an aberrant phenotype was visually apparent at 6 hours post onset of streak regression or earlier. We performed PIV on the remaining $N_{\text{healthy}} = 71$ embryos to determine the time point of onset of streak regression as explained below. In 9 embryos, an onset of streak regression could not be determined, because the streak regresses already at the first time frame or is not regressing yet at the last time frame. Only the remaining $N_{\text{with } t_0} = 63$ embryos were used for further analysis.

| Condition | $N_{\text{total}}$ | $N_{\text{healthy}}$ | $N_{\text{with } t_0}$ |
|---|---|---|---|
| Control | 28 | 26 | 25 |
| ventral tissue removed | 25 | 18 | 16 |
| ventral tissue replaced | 22 | 17 | 13 |
| dorsal vitelline removed | 14 | 10 | 9 |

**Table S1:** Number of embryos per condition.

##### 1.2.2 Particle image velocimetry

We used PIVlab [68] to infer velocities of 32x32 pixel windows corresponding to a grid of $20\mu$mx$20\mu$m overlapping squares with the centers of neighboring squares separated by $10\mu$m. Three passes were used with window widths of 128, 64, 32 pixels. We applied an absolute threshold of $5\mu$m/min to the velocities, followed by a normalized median test [69], as implemented in PIVlab, using $\varepsilon = 3\mu$m/min and a normalized threshold of 3. The thus excluded data points were not interpolated.

##### 1.2.3 Aligning embryos in space and time

In order to spatially align the data from different embryos, the position of the Hensen's node and the position and orientation of the mid-line defined by the primitive streak were manually annotated. To define the mid-line, we annotate two points $\mathbf{r}_1$, $\mathbf{r}_2$ on the center-line of the primitive streak (see Fig. S3a for an example). We identified the contour of the streak based on the cell debris that accumulate at the middle of the streak and yield a bright signal in the membrane-GFP channel. The center of the node is also identified based on the elongated morphology of the surrounding cells. The two points $\mathbf{r}_1$, $\mathbf{r}_2$ define the orientation of the mid-line, i.e. a normalized vector parallel to the mid-line defined as

$$\mathbf{n} = (\mathbf{r}_1 - \mathbf{r}_2)/|\mathbf{r}_1 - \mathbf{r}_2|. \tag{S1}$$

To define the position of the mid-line, we project the annotated position of the node $\mathbf{r}_{\text{n}}$ onto the mid-line:

$$\mathbf{r}_{\text{n,mid}} = \mathbf{r}_1 + [\mathbf{n} \cdot (\mathbf{r}_{\text{n}} - \mathbf{r}_1)]\,\mathbf{n} \tag{S2}$$

We manually annotate node and streak at time-points separated by 20min. We then interpolate the position of the node and the orientation vector $\mathbf{n}$ at the remaining time-points of the experiment using a Gaussian kernel with a width of 20min. Thus, we obtain a reference frame defined by $\mathbf{r}_{\mathrm{n,mid}}$ and $\mathbf{n}$ at each time-point of an experiment.

With this, we interpolate the flow field at grid points of a square grid with a width of 2000$\mu$m centered at $\mathbf{r}_{\mathrm{n,mid}}$ identifying the axis of the streak (i.e. the AP axis and mid-line of the embryo) as the $y$ axis. We use a grid spacing of 25$\mu$m and a Gaussian kernel with a width $\sigma = 25\mu$m for interpolation. Using this grid, we average the flow field of each embryo in 1h time windows around the original time points of the experiment. From this, we determine the anterior-posterior velocity $v_y$ of the node with respect to the surrounding epiblast, by calculating the average flow field within a radius of 100$\mu$m around the node and subtracting the average flow field in rectangles with coordinates $|x| \in [200\mu\mathrm{m}, 400\mu\mathrm{m}]$ and $y \in [-300\mu\mathrm{m}, 300\mu\mathrm{m}]$ with respect to the node. In Fig. S2a this velocity is plotted for 26 unperturbed embryos as function of time $t_{\mathrm{exp}}$ since the start of the experiment. The velocity $v_y$ is positive for early time points, as the streak elongates and the node moves anteriorly. At late times, $v_y$ is negative as the streak regresses and the node moves posteriorly. We use the last time point where $v_y > 0$ as the reference time point $t = 0$ corresponding to the onset of streak regression. In Fig. S2b, $v_y$ is plotted as a function of $t$ for $N_{\mathrm{with}\ t_0} = 25$ embryos. We observe that the data from all experiments collapse onto a common curve, which validates our method for spatial and temporal alignment. This allows us to compare flow fields at a given point in space and time. To this end, we use the grid with respect to the mid-line as described above and average for each embryo the flow field in a 1h time window around time points separated by $0.5h$ starting from $t = -10h$. In the main text, we focus on the time point $t = 0$.

##### 1.2.4 Flow field decomposition and median flow calculation

We observe that the tissue around the streak often translates or rotates as a whole with respect to the lab frame. As we focus here on the flow at the node, we subtract such embryo-scale rigid body components from the flow. To this end, we take into account data points with a distance between 350$\mu$m and 600$\mu$m with respect to $\mathbf{r}_{\mathrm{n,mid}}$. Using these data points we calculate a rigid body rotation rate $\Omega$ and a velocity $\mathbf{V}$ as

$$\mathbf{V} = \langle \mathbf{v} \rangle\,, \quad \Omega = \mathbf{z} \cdot \langle \mathbf{r} \times \mathbf{v} \rangle \,/\, \left\langle |\mathbf{r}|^2 \right\rangle \tag{S3}$$

We find that the left-right component of $\mathbf{V}$ and the rotation rate $\Omega$ do not have a consistent sign across embryos (Fig S1). Hence they do not amount to a chiral flow that could facilitate the definition of a left-right axis. However, the magnitude of $\mathbf{V}$ in single embryos is often on the order of magnitude of the chiral tissue flow at the node, which complicates the quantification of chiral flow. Therefore, we focus in the following only on the residual of the measured flow field after subtracting the rigid body motion of the tissue surrounding the node, i.e. the node frame velocity

$$\mathbf{v}_{\text{node-frame}} = \mathbf{v} - \mathbf{V} - \Omega \mathbf{z} \times \mathbf{r}. \tag{S4}$$

Then, we decompose the flow field $\mathbf{v}_{\text{node-frame}}$ into a left-right symmetric and a left-right anti-symmetric component according to

$$\mathbf{v}_{\text{sym}}(x,y) = \frac{1}{2}\left(v_x(x,y) - v_x(-x,y), v_y(x,y) + v_y(-x,y)\right)^T \tag{S5}$$

$$\mathbf{v}_{\text{antisym}}(x,y) = \frac{1}{2}\left(v_x(x,y) + v_x(-x,y), v_y(x,y) - v_y(-x,y)\right)^T. \tag{S6}$$

Then, we calculate median flow fields across the set of 25 embryos, by calculating the median components $(v_x, v_y)$ at each point in space and time. We calculate this median for the symmetric component, antisymmetric component and for the total flow field $\mathbf{v}_{\text{node-frame}}$ separately. We also quantify the embryo-to-embryo variability in terms of the standard deviation at all points in space and time and for all components of the flow field. We find that the magnitude of the median left-right antisymmetric component is less than the standard deviation everywhere except at the node for $t \leq 0$ (Fig. S4).

##### 1.2.5 Quantification of flow in single embryos

In order to have a metric that allows for a quantitative comparison of chiral flow among embryos under different conditions, we quantify the rotation of the node relative to the surrounding tissue. To this end, we use the position $\mathbf{r}_\text{n}$ of the node center that we annotated manually as described above. We calculate the flow field on a grid as before, but with the origin of the grid defined by $\mathbf{r}_\text{n}$ instead of $\mathbf{r}_{\text{n,mid}}$. From this we calculate the rotation speed of the node relative to the surrounding tissue as

$$\Omega_{\text{node,rel}} = \frac{\mathbf{z} \cdot \langle \mathbf{r} \times \mathbf{v} \rangle_{|\mathbf{r}_i - \mathbf{r}_\text{n}| \leq 50\mu\text{m}}}{\langle |\mathbf{r}|^2 \rangle_{|\mathbf{r}_i - \mathbf{r}_\text{n}| \leq 50\mu\text{m}}} - \frac{\mathbf{z} \cdot \langle \mathbf{r} \times \mathbf{v} \rangle_{75\mu\text{m} < |\mathbf{r}_i - \mathbf{r}_\text{n}| \leq 400\mu\text{m}}}{\langle |\mathbf{r}|^2 \rangle_{75\mu\text{m} < |\mathbf{r}_i - \mathbf{r}_\text{n}| \leq 400\mu\text{m}}} \tag{S7}$$

We do so for each embryo and each time frame of an experiment and calculate the average in 1h time-windows for each embryo and time-points separated by 0.5h with respect to the onset of streak regression ($t = 0$). Analogously we calculate the left-right velocity of the node as

$$V_{\text{LR,rel}} = \langle v_x \rangle_{|\mathbf{r}_i - \mathbf{r}_\text{n}| \leq 50\mu\text{m}} - \langle v_x \rangle_{75\mu\text{m} < |\mathbf{r}_i - \mathbf{r}_\text{n}| \leq 400\mu\text{m}}. \tag{S8}$$

It has recently been claimed that there are consistent left-right asymmetries in the large-scale polonaise tissue flows preceding the chiral flow at the Hensen's node [70]. In order to quantify polonaise flows and their left-right asymmetry we quantified the average vorticity in the left and the right half of the embryo, again using the flow field in a grid centered at $\mathbf{r}_{\text{n,mid}}$:

$$\Omega_L = \langle \text{rot } \mathbf{v} \rangle_{|\mathbf{r}| < 600\mu\text{m} \wedge x < -100\mu\text{m}}, \quad \Omega_R = \langle \text{rot } \mathbf{v} \rangle_{|\mathbf{r}| < 600\mu\text{m} \wedge x > 100\mu\text{m}} \tag{S9}$$

$\Omega_L - \Omega_R$ is a measure of the left-right symmetric polonaise flow that accompany streak extension, whereas during streak regression a vortex pair with opposite handedness arises (Fig. S1d). $\Omega_L + \Omega_R$ quantifies the left-right asymmetry of the polonaise flow. We find no consistent asymmetry at any time point. We note, however, that we have only a few data points at early times ($t < -5h$), though still more than in [70]. Furthermore, we restrict our analysis to the vicinity of the node ($|\mathbf{r}| < 600\mu$m), because further away we have too few data points.

In addition, we quantified the flux density $J_{\text{streak}}$ of material flowing into the streak as the integrated divergence at the streak and the node

$$J_{\text{streak}} = \int_{|\mathbf{r}|<100\mu\text{m} \vee (|x|<100\mu\text{m} \wedge 600\mu\text{m}<|y|<0} \mathrm{d}S \,\text{div}\,\mathbf{v} \tag{S10}$$

We use here the flow field in a grid centered at $\mathbf{r}_{\text{n,mid}}$ and use finite differences to calculate the divergence. We find that the flux into the streak peaks before streak regression. In mechanically perturbed embryos, the quantification of $J_{\text{streak}}$ shows that cells continue to ingress at similar speed when compared with unperturbed embryos, which supports the notion that the flux towards the streak is mechanically driven and controlled within the epiblast (Fig. S5i-l).

For all those measures we calculate the median across embryos. Furthermore, we determine for each time point whether the median is different from zero at the 1% significance level using a Wilcoxon signed rank test as implemented in MATLAB [61].

##### 1.2.6 Determining the onset of chiral flow and integrated node rotation

In order to determine the onset of chiral flow, we consider the node rotation $\Omega_{\text{node,rel}}$ calculated as described above (Eq. S7). In each embryo, we calculate the time point $t_{\max}$ where the 1h moving average of $\Omega_{\text{node,rel}}$ is maximal. We define the onset of chiral flow in terms of the half-maximum, i.e. as the time point $t_{\text{chir}}$ of the experiment where the 1h average $\Omega_{\text{node,rel}}$ is less than half the maximum value $\Omega_{\text{node,rel}}$ at $t_{\max}$ for the first time when going back in time from the time point $t_{\max}$. In Fig. 3j, we plot $t_{\text{chir}}$ with respect to the onset of streak regression at $t = 0$.

We use the thus determined onset of chiral flow to calculate an integrated node rotation

$$\theta_{\text{flow}} = \int_{t_{\text{chir}}-1h}^{t_{\text{chir}}+5h} \mathrm{d}t\, \Omega_{\text{node,rel}}(t), \tag{S11}$$

where again $\Omega_{\text{node,rel}}$ is calculated from the flow field (Eq. S7) and averaged in 1h time windows before calculating the integral.

##### 1.2.7 Quantification of kink angle

We quantify also the left-right asymmetric morphology of the node. We observe that during streak regression the most anterior part of the streak kinks to the left. We quantify this kink using manual annotation: We annotate two points that define the mid-line of the streak. Then we annotate two points that define the long axis of the node (See Fig S3b for an example). The node is identified based on the accumulation of cell debris at its center and the characteristic elongated morphology of the surrounding cells. From these two lines, we calculate the kinking angle as the angle between these lines. Manual annotation was done by two authors independently and the mean angle between the two annotation was used. We then determined the maximum kink angle for time points $-4h \leq t < 0h$ and $0h < t \leq 6h$, as plotted in Fig. 1k.

##### 1.2.8 Analysis of laser ablation experiments

From the time-lapse microcopy images, we determined the start of the laser ablation. The ablation itself takes about 1-2$s$ (See mov2). For quantifying the elastic response of the tissue, we compare the last frame before the laser ablation with the frame 5$s$ after the start of the laser ablation. As for the long term tissue flows, we used PIVlab [68] to infer the displacement $\mathbf{u}$ of the tissue between these frames. As before we used three passes with window widths of 128, 64 and 32 pixels. We removed outliers using a normalized median test with $\varepsilon = 3\text{px}$ and a normalized threshold of 3 [69].
Using the frame 5s after the start of the laser ablation, we manually annotated 10 points marking the outline of the lasercuts. From this, we determined a mask of the cut using the `poly2mask` function of MATLAB [61]. From this mask, we calculate the center of mass position $\mathbf{r}_{\text{mid}}$ of the cut side.
In order to quantify the rotation of the ablated tissue, we calculated a rotation angle $\beta_{\text{in}}$ of a 50$\mu$m circle around $\mathbf{r}_{\text{mid}}$ and a rotation $\beta_{\text{out}}$ of a ring surrounding the cut side analogous Eq. S7 as

$$\beta_{\text{in}} = \frac{\mathbf{z} \cdot \langle \mathbf{r} \times \mathbf{u} \rangle_{|\mathbf{r}_i - \mathbf{r}_{\text{mid}}| \leq 50\mu\text{m}}}{\langle |\mathbf{r}|^2 \rangle_{|\mathbf{r}_i - \mathbf{r}_{\text{mid}}| \leq 50\mu\text{m}}}, \quad \beta_{\text{out}} = \frac{\mathbf{z} \cdot \langle (\mathbf{r}_i - \mathbf{r}_{\text{mid}}) \times \mathbf{u}_i \rangle_{125\mu\text{m} < |\mathbf{r}_i - \mathbf{r}_{\text{mid}}| \leq 200\mu\text{m}}}{\langle |\mathbf{r}_i - \mathbf{r}_{\text{mid}}|^2 \rangle_{125\mu\text{m} < |\mathbf{r}_i - \mathbf{r}_{\text{mid}}| \leq 200\mu\text{m}}} \tag{S12}$$

Note that we use circular cuts of varying sizes, with a radius of up to $50 - 150\mu$m. In any case, we exclude data points within the mask of the cut side from the calculation of $\theta_{\text{out}}$. Analogously, we also quantified the net leftward displacement of the tissue upon circular laser cuts:

$$x_{\text{in}} = \langle \mathbf{u} \cdot \mathbf{x} \rangle_{|\mathbf{r}_i - \mathbf{r}_{\text{mid}}| \leq 50\mu\text{m}}, \quad x_{\text{out}} = \langle \mathbf{u} \cdot \mathbf{x} \rangle_{125\mu\text{m} < |\mathbf{r}_i - \mathbf{r}_{\text{mid}}| \leq 200\mu\text{m}}. \tag{S13}$$

We find no evidence for a net displacement (Fig. S11), indicating that the node does not generate a net force with respect to the underlying tissue substrate.

##### 1.2.9 Bootstrapping

In order to obtain a measure of the experimental uncertainty, we made use of bootstrapping. Using random sampling with replacement, we generated 400 sets of $N_{\text{healthy}}$ embryos from the original data set for each experimental condition. For each set of embryos, we calculated median quantities, i.e. the median across embryos of e.g. the rotation speed $\Omega_{\text{node,rel}}$. From this set of median quantities, the $5^{\text{th}}$ and $95^{\text{th}}$ percentiles were determined. This defines the range of uncertainty of the median, which we give in the main text.

### 2 Supplementary notes on physical theory

#### 2.1 Fluid model of the epiblast

In the following, we briefly introduce a generic mechanical model of large-scale tissue flows in the early quail embryo. In subsection 2.3, we use this model to infer an active force density that drives the flows during left-right symmetry breaking, as discussed in the main text.

The epiblast is a rapidly proliferating epithelial tissue. In such a tissue, the continuous turnover facilitates the relaxation of elastic stresses thereby fluidising the tissue [72]. Motivated by this, we adopt a fluid model to capture the experimentally observed large-scale tissue flows in the epiblast, similar to previous approaches [52,44,53]. For simplicity, we consider a flat fluid film neglecting the cell-scale curvature of the epiblast surface close to the streak and the node.

We consider a shear viscosity $\eta$ and a bulk viscosity $\alpha\eta$ where $\alpha$ is a dimensionless parameter. With this the viscous stress reads

$$t_{ij}^{\text{visc}} = \eta\left[2v_{ij} + \delta_{ij}(\alpha - 1)\text{div}\,\mathbf{v}\right], \tag{S14}$$

where $v_{ij} = 1/2(\partial_i v_j + \partial_j v_i)$ is the shear rate tensor associated with tissue flow field $\mathbf{v} = v_x\mathbf{x} + v_y\mathbf{y}$. Mechanical activity of the tissue—such as due to actomyosin cables as well as proliferation—give rise to an additional contribution $t_{ij}^{\text{act}}$ to the stress tensor of the epiblast:

$$t_{ij} = t_{ij}^{\text{visc}} + t_{ij}^{\text{act}}, \tag{S15}$$

The symmetric component of the stress tensor captures mechanical interaction within the epiblast, including both viscous and active nematic or isotropic stresses. In a chiral active fluid, mechanical activity can give rise to an antisymmetric active stress [23,24,25,50]. Importantly, such an antisymmetric stress corresponds to a torque density

$$\tau = \epsilon_{ji} t_{ij}^{\text{act}}, \tag{S16}$$

where $\epsilon_{ij}$ is the Levi-Civita-Tensor and we adopt Einstein sum convention over repeated indices

[50,73]. Locally, such torques acting on the epiblast tissue can result from torque dipoles in the plane. We denote the density of such in-plane torque dipoles as the moment tensor $M_i^{\mathrm{act}}$, i.e. a pseudo-vector field corresponding to the tangential transport of the $z$ component of angular momentum [50,73,23,24]. In-plane torque dipoles drive rotations of some segment of the epiblast relative to the surrounding epiblast. As such, they cannot yield a net torque exerted on the epiblast. In contrast, such a torque can result from mechanical coupling of the epiblast to an underlying substrate. Active chiral processes at the interface of the epiblast and the substrate, i.e. the meso/endoderm tissue, can yield microscopic torque-dipoles. On a hydrodynamic scale, these torque dipoles result in a torque density $\tau_{\mathrm{act}}$ that the substrate exerts on the epiblast. With this, the torque balance equation for the epiblast reads

$$\epsilon_{ji} t_{ij}^{\mathrm{act}} = \tau_{\mathrm{act}} + \partial_i M_i^{\mathrm{act}}. \tag{S17}$$

Mechanical coupling to a rigid substrate also yields a friction force density $-\gamma \mathbf{v}$, which defines a hydrodynamic length scale $l_{\mathrm{hyd}} = \sqrt{\eta/\gamma}$ (for more detailed discussion of modeling the mechanical coupling of tissue layers see section 2.6). For completeness, we also consider an active force density $\mathbf{f}_{\mathrm{sub,act}}$ that propels the epiblast relative to the substrate. Then, tangential force balance gives the governing equation of the epiblast flow field

$$\partial_j (t_{ji}^{\mathrm{act}} + t_{ji}^{\mathrm{visc}}) = \gamma v_i - f_i^{\mathrm{sub,act}}. \tag{S18}$$

In the experiment, only the flow field and its derivative are accessible. This motivates us to rewrite the force balance equation as

$$-\partial_j t_{ji}^{\mathrm{visc}} + \gamma v_i = f_i^{\mathrm{act}} \tag{S19}$$

where the active force density $\mathbf{f}_{\mathrm{act}}$ is defined as

$$f_{\mathrm{act}}^i = f_{\mathrm{act,sub}}^i + \partial_j t_{ji}^{\mathrm{act}} \tag{S20}$$

Two patterns of active stresses and substrate forces that result in an identical active force density $\mathbf{f}_{\mathrm{act}}$ yield identical flow fields. Thus, the pattern of active stresses cannot be uniquely inferred from the observed flow field, only $\mathbf{f}_{\mathrm{act}}$ can be inferred, given that we know the viscosities and the hydrodynamic length of the tissue (see also section 2.3).

#### 2.2 Inferring material parameters of the epiblast

Away from the streak, the epiblast appears homogeneous in terms of tissue architecture prior to streak regression. Thus, we hypothesize that material properties are spatially homogeneous, i.e. viscosities and hydrodynamic length are constant throughout the epiblast except at the streak and the node. Furthermore, we hypothesize that also mechanical activity away from the streak is constant and isotropic such that $t_{ij}^{\mathrm{act}}$ = const. and thus $\mathbf{f}_{\mathrm{act}} = 0$, To test this hypothesis, we consider a boundary that encompasses the epiblast in a distance of up to 600$\mu$m of the node and excludes the tissue surrounding the node and the streak in a distance of up to 100$\mu$m (see Fig S6e,l). If the tissue enclosed in this boundary behaves as a homogeneous fluid film, the flow field in this region can be calculated from velocities at the boundary, as we describe in the following.

##### 2.2.1 Analytic solutions for constant material parameters

For $\mathbf{f}_{\text{act}} = 0$, the force balance equation (Eq. S19) yields the following differential equation governing the flow field:

$$\Delta v_i + \alpha\partial_i(\partial_j v_j) - \frac{1}{l_h^2} v_i = 0, \tag{S21}$$

where $\Delta$ denotes the Laplace operator in Cartesian coordinates. We write the flow field as a Hodge decomposition

$$v_i = \partial_i A + \epsilon_{ji}\partial_j B, \tag{S22}$$

where $A$ and $B$ correspond to irrotational and rotational components of the flow field [74,75,76]. We plug this into the force balance equation and find that it yields independent equations for the irrotational and rotational flow components:

$$(1+\alpha)\Delta A - \frac{1}{l_h^2} A = 0 \tag{S23}$$

$$\Delta B - \frac{1}{l_h^2} B = 0 \tag{S24}$$

Let us consider an annulus, i.e. a domain $\{\mathbf{r} \in \mathbb{R}^2 \big| a < |\mathbf{r}| < R\}$ with $a, R > 0$. Using polar coordinates $r, \theta$ solutions are given by a multipole expansion:

$$A = \text{Re}\left\{\sum_{m=0}^{\infty}\left[A_m^- K_{|m|}\left(\frac{r}{l_h^A}\right) + A_m^+ I_{|m|}\left(\frac{r}{l_h^A}\right)\right] e^{im\theta}\right\}, \tag{S25}$$

$$B = \text{Re}\left\{\sum_{m=0}^{\infty}\left[B_m^- K_{|m|}\left(\frac{r}{l_h^B}\right) + B_m^+ I_{|m|}\left(\frac{r}{l_h^B}\right)\right] e^{im\theta}\right\}, \tag{S26}$$

where $A_m^{+/-}$ and $B_m^{+/-}$ are complex coefficients and the hydrodynamic lengths are defined as $l_h^B = l_h$, $l_h^A = \sqrt{\alpha+1} l_h$. The modified Bessel functions are defined as

$$I_m(r) = \lim_{\alpha\to m}\sum_{n=0}^{\infty}\frac{1}{n!\Gamma(\alpha+n+1)}\left(\frac{r}{2}\right)^{2n+\alpha}, \tag{S27}$$

$$K_m(r) = \lim_{\alpha\to m}\frac{\pi}{2}\frac{I_{-\alpha}(r) - I_\alpha(r)}{\sin\alpha\pi}, \tag{S28}$$

with $\Gamma(n)$ denoting here the gamma function. $I_{|m|}(r)$ is monotonically increasing with $r$ whereas $K_{|m|}(r)$ is monotonically decreasing. For $a \ll R$, $A_m^-, B_M^-$ correspond to Fourier components of the boundary velocities or forces at the inner ring, i.e. $|\mathbf{r}| = a$, whereas $A_m^+, B_m^+$ represent the boundary conditions at the outer ring, $|\mathbf{r}| = R$. This expansion allows to discuss a model where the epiblast is a homogeneous material everywhere but at the node and in a far distance from the node. To this end, we consider an annulus centered at the node with $a = 100\mu$m and $R = 600\mu$m.

However, we want to also allow for distinct material properties, in particular mechanical activity, of the streak. To this end, we consider a density of forces and force dipoles localized to the center of the streak. The flow field resulting from a force monopole, i.e. a force density $\mathbf{F}\delta^{(2)}(x-x', y-y')$ at position $(x', y')$ in an infinite plane can be written as

$$v_y - iv_x = G_{\text{mono}}(\mathbf{r}-\mathbf{r}', \mathbf{F}) = G_0(\mathbf{r}-\mathbf{r}')(F_y - iF_x) + G_2(\mathbf{r}-\mathbf{r}')(F_y - iF_x), \tag{S29}$$

where the complex-valued Green's functions (see also [77]) are defined as

$$G_0(\mathbf{r}) = \frac{1}{4\pi\eta}\left[\frac{1}{\alpha+1}K_0(|\mathbf{r}|/l_h^A) + K_0(|\mathbf{r}|/l_h)\right] \tag{S30}$$

$$G_2(\mathbf{r}) = \frac{1}{4\pi\eta}\left[\frac{1}{\alpha+1}K_2(|\mathbf{r}|/l_h^A) - K_2(|\mathbf{r}|/l_h)\right]\frac{(y-ix)^2}{|\mathbf{r}|^2}. \tag{S31}$$

A force dipole, given by a pair of forces $\pm\mathbf{F}$ separated by $d\boldsymbol{\nu}$ centered at $\mathbf{r}'$, yields a flow field

$$v_y - iv_x = G_{\text{dip}}(\mathbf{r}, \mathbf{r}', \mathbf{F}, d\boldsymbol{\nu}) = d(\nu_i\partial_i)G_{\text{mono}}(\mathbf{r}, \mathbf{r}', \mathbf{F}). \tag{S32}$$

Taken together, this allows us to write the flow field in an annulus with additional forces and force dipoles localized to a line, not unlike a crack.

##### 2.2.2 Calculating flow field from measured boundary velocities

To analyze experimental data, we consider forces and force dipoles at $N_C$ discrete points along the streak. Furthermore, we cut off the multipole expansion (corresponding to force multipoles at $r \to 0$ and $r \to \infty$) at $N_-$ and $N_+$, respectively, yielding

$$\begin{aligned} v_y - iv_x = &\sum_{m=0}^{N_+}\left[a_m^+ V_m^{A,+,r}(z) + ib_m^+ V_m^{A,+,i}(z) + c_m^+ V_m^{B,+,i}(z) + id_m^+ V_m^{B,+,r}(z)\right] \\ &+ \sum_{m=0}^{N_-}\left[a_m^- V_m^{A,-,r}(z) + ib_m^- V_m^{A,-,i}(z) + c_m^- V_m^{B,-,i}(z) + id_m^- V_m^{B,-,r}(z)\right] \\ &+ \sum_{j=1}^{N_C}\big[A_j G_{\text{mono}}(\mathbf{r}-\mathbf{r}_j, \mathbf{y}) + B_j G_{\text{mono}}(\mathbf{r}-\mathbf{r}_j, \mathbf{x}) \\ &\qquad + D_j G_{\text{dipo}}(\mathbf{r}-\mathbf{r}_j, \mathbf{y}, \mathbf{x}) + C_j G_{\text{dipo}}(\mathbf{r}-\mathbf{r}_j, \mathbf{x}, \mathbf{x})\big], \end{aligned} \tag{S33}$$

where

$$V_m^{A/B,-,r}(z) = K_{m+1}(r/l_h^{A/B})e^{i(m+1)\theta} + K_{m-1}(r/l_h^{A/B})e^{-i(m-1)\theta} \quad \text{(S34)}$$

$$V_m^{A/B,-,i}(z) = K_{m+1}(r/l_h^{A/B})e^{i(m+1)\theta} - K_{m-1}(r/l_h^{A/B})e^{-i(m-1)\theta} \quad \text{(S35)}$$

$$V_m^{A/B,+,r}(z) = I_{m+1}(r/l_h^{A/B})e^{i(m+1)\theta} + I_{m-1}(r/l_h^{A/B})e^{-i(m-1)\theta} \quad \text{(S36)}$$

$$V_m^{A/B,+,i}(z) = I_{m+1}(r/l_h^{A/B})e^{i(m+1)\theta} - I_{m-1}(r/l_h^{A/B})e^{-i(m-1)\theta} \quad \text{(S37)}$$

and $\mathbf{x}, \mathbf{y}$ denote the unit basis vectors of the Cartesian coordinate system with the mid-line of the embryo coinciding with the $y$-axis ($\theta \in \{0, \pm\pi\}$). $a_m^{+/-}$, $b_m^{+/-}$, $c_m^{+/-}$, $d_m^{+/-}$, $A_j$, $B_j$, $D_j$, $C_j$ are real coefficients, which we determine from the measured boundary velocities using linear regression. $a_m^{+/-}$, $c_m^{+/-}$, $A_j$ and $C_j$ yield the left-right symmetric flow field, where as $b_m^{+/-}$, $d_m^{+/-}$, $B_j$ and $D_j$ yield the left-right antisymmetric flow field.

##### 2.2.3 Analysis of symmetric flow field

First, we focus on the left-right-symmetric tissue flow towards the streak. Previous studies of earlier stages have shown that the flow towards the streak can be understood in terms of contractile mechanical activity localized to the streak [44,52,53]. As a proof of principle, we ask if we do indeed need to consider mechanical activity at the streak to understand the flow field. To this end, we consider as an outer boundary a circle with radius 600$\mu$m centered at the node. We linearly interpolate the measured left-right symmetric flow field at $(2N_+ + 1)$ equally distant points along this circle, where we use $N_+ = 63$. This defines a velocity boundary condition. To calculate the resulting flow field inside the circle for given $l_h$ and $\alpha$, we calculate the $(2N_+ + 1)$ coefficients $a_m^+$, $c_m^+$ from the $(2N_+ + 1)$ boundary velocities using Eq. S33.

We consider a wide range of parameters (Fig. S6o). For each pair of parameters we calculate the residual of the experimental flow field after subtracting the flow field given by Eq. S33. We calculate the root mean squared of the residual for data points inside the 600$\mu$m circle and time points $t \in \{-3h, -2h, -1h, 0h\}$ relative to streak regression. We find that the experimental flow field is best described for large hydrodynamic length and $\alpha \sim 1$. However, even when using these optimal parameters the calculated flow field does not capture the flow towards the streak. Thus, the flow towards the streak does indeed imply that the streak is mechanically distinct from the surrounding epiblast.

Next we ask, if taking into account forces and force dipoles generated at the streak is sufficient to capture the large-scale flow field including the flow towards the streak. To this end we use data points for which the minimal distance $r_{\min}$ to the negative y axis obeys $100\mu\text{m} \leq r_{\min} < 125\mu\text{m}$ and for which $r < 600\mu\text{m}$. This yields on each side of the streak $2N_C$ data points with $y < 0$, i.e. anterior to the node center with $N_C = 20$. Furthermore, we have $2(2N_- + 1) + 1$ data points with $x \geq 0$ (i.e. posterior to the node center) , where $N_- = 4$. With this and the 127 equally distant points around the bounding circle, we calculate the coefficients $a_m^{+/-}$, $c_m^{+/-}$, $A_j$ and $C_j$. As before we determine the root mean square residual of the experimental flow field for a range of material parameters $\alpha, l_h$ (Fig. S6p). When considering time points up to the onset of streak regression, the calculated flow field is in very good agreement with the experimental flow field for a large hydrodynamic length $l_h \geq 300\mu$m. We conclude that the large-scale tissue flows up to the time point of streak ingression onset are well captured by a fluid model with mechanical activity localized to the streak and the node. Notably, there is no signature of a finite hydrodynamic length, indicating that mechanical coupling to a substrate, in particular the vitelline membrane, is negligible (see the section 2.6 for a more detailed discussion).

##### 2.2.4 Analysis of chiral flow

Analogously, we analyzed the left-right antisymmetric component of the median flow field (Fig. S7). Again, we consider time points from the onset of chiral flow till the onset of streak regression, i.e. $t \in \{-3h, -2h, -1h, 0h\}$. We find that the flow field calculated from the velocities at the boundary of streak and node captures the experimental flow field in the surrounding well for a hydrodynamic length of about $75\mu$m and $\alpha \sim 1$ (Fig. S7k) Only for such a small hydrodynamic length does the calculated flow field capture the sharp decay of chiral flow away from the node (Fig. S7l,m). This implies that the mechanical nature of chiral flow is distinct from the large-scale flow towards for the streak, as for the latter no finite hydrodynamic length is evident. We discuss this observation further in section 2.6. We conclude that the measured spatial profile of chiral flow is consistent with a fluid model, where chiral flow is driven by mechanical activity localized to the node and the streak, and limited by mechanical coupling to a substrate.

##### 2.2.5 Estimating the viscosity of the epiblast

While, $\alpha$ and $l_h$ can be estimated from flow profiles, the absolute value of the viscosity cannot be inferred without measuring forces. Until very recently (see [55]), no measurements of the 2D viscosity of an epithelium subject to in-plane deformations have been reported. However, we can obtain a rough estimate based on literature values of viscosity and elasticity in different tissues. These values differ by several orders of magnitude, also due to differences in the considered time and length scales. Still this values inform a central estimate, which is in remarkable agreement with measurements in the avian epiblast that were published while preparing this manuscript [55].

A particularly elegant method to measure the viscosities of embryonic tissue *in vivo* is based on ferrofluidic droplets that are deforming due to an externally applied magnetic field [78]. Inserting droplets of radius $20-40\mu$m into bulk tissue of zebra fish embryos, a viscosity of $\eta_{3D} \sim 5\,$kPa s has been measured. However, we note that in these experiments the tissue was subject to only subtle deformations involving little to no cell rearrangements. This is also evident from the relaxation time scale $t_r \sim 1$min, which corresponds to the relaxation time scale of the cytoskeleton [79,80].

Thus, we suspect that this method yields a measure of cell-internal viscosity, which may differ by several orders of magnitude from the tissue viscosity relevant to large-scale morphogenetic flows.
An alternative, though significantly more disruptive, method is micropette aspiration [81]. Applied to 3D cell aggregates using a pipette radius of $35\mu$m, a viscosity $\eta_{3D} = 200\,$kPa s and a relaxation time scale $t_r = 45$min corresponding to an elastic modulus $E_{3\mathrm{D}} = 700$Pa have been obtained [81]. Similar values have been obtained by compressing spherical cell aggregates several 100 $\mu$m in size: $\eta_{3D} = 440$kPa s, $t_r = 5$h, $E_{3D} = 25$Pa [82]. Thus, we estimate the 3D viscosity of a bulk tissue as $\eta_{3D} \sim 100 - 1000$kPas. Multiplying with the thickness of about $50\mu$m of the avian epiblast, we obtain a 2D viscosity of $\eta \sim 5 - 50$Ns/m.

However, we expect that the mechanical properties of an epithelium differ considerably from a thin layer of bulk tissue. In particular, stresses may be concentrated in the apical layer, which may even make the mechanical properties of an epithelium largely independent of its thickness. Mechanical properties of embryonic epithelia have been obtained for the *Xenopus laevis* gastrula [83] and the sea urchin blastula [84]. Using compression of the sea urchin blastocoel wall, a 2D elasticity of $E_{2D} \sim 10\mathrm{mN}/m$ and a 2D viscosity of $\eta \sim 0.1-1Ns/m$ have been observed [84]. For isolated epithelia from *Xenopus laevis*, mechanical properties have been estimated by analyzing the shape as the epithelium is deformed by gravity, yielding $E_{2D} = 1$mN/m, $\eta = 0.4Ns/m$ for a relaxation time scale of $t_r = 5$min. This time scale is considerably smaller than what we would expect from tissue fluidized by cell divisions [72]. Furthermore, we note than in both experiments, the response of an epithelial surface to 3D deformations was analyzed, which may not yield a reliable estimate for the viscous response to in-plane shear deformations. A more reliable measurement of 2D mechanical properties may be obtained from stretching a freely suspended epithelial monolayer in the plane as was done in [80] using an MDCK cell culture. This yields a 2D elastic modulus $E_{2D} = 200$mN/m. Using a relaxation time scale $t_r > 1$min, this yields a 2D viscosity $\eta_{2D} > 10Ns/m$.

Taken together we obtain estimates of $\eta \sim 0.1 - 100$Ns/m, yielding a central estimate of $\eta \approx 3\mathrm{Ns}/m \approx 1\mathrm{mN\,h/m}$, which is in remarkable agreement with the measured value ($\eta = 4\mu\mathrm{Ns}/\mu\mathrm{m} \sim 1\mathrm{mN\,h/m}$) in [55] . This yields active force densities (Fig. 2a-d) on the order of $|\mathbf{f}_{\mathrm{act}}| \sim 10$Pa, which is consistent with measured tangential traction forces ($5 - 20$Pa) from traction force microscopy of epithelial tissues [85].

#### 2.3 Inferring the forces driving epiblast flow

So far, we have established that the streak and the node are, as a material, mechanically distinct from the surrounding epiblast. Next, we set out to characterize the streak tissue in terms of the forces it generates to drive the tissue flows in the surrounding epiblast. In particular, we calculate the active force density $\mathbf{f}_{\mathrm{act}}$ (Eq. S19). We use the material parameters $l_h, \alpha$ determined in the previous section. We note that we expect the viscosities of the streak to be distinct from the surrounding epiblast. As such, the force field we calculate from Eq. S19 does not yield the true active forces $\mathbf{f}_{\text{act}}$, but it yields an effective picture of what the streak does to the epiblast in terms of forces. Specifically, integrating $\mathbf{f}_{\text{act}}$ across the width of the streak yields a line density of forces and force-dipoles that the tissue of the streak exerts onto the epiblast. Furthermore, integrating $\mathbf{f}_{\text{act}}$ over the node area yields a net force and torque exerted on the surrounding epiblast.

Calculating derivatives in terms of finite differences of the median flow field yields noisy results. Instead, we calculate the (second order) derivatives in Eq. S19 for each embryo separately using finite differences applied to the 1h moving average of the measured flow field. We then smoothen the calculated field $\mathbf{f}_{\text{act}}$ using a Gaussian kernel with $\sigma = 25\mu\text{m}$. Subsequently, we calculate the median $\mathbf{f}_{\text{act}}$ across all 25 embryos. Finally, we apply once a again Gaussian smoothening with $\sigma = 25\mu\text{m}$.

As before, we analyze the left-right symmetric and antisymmetric components of the median flow field separately. We focus on the time point where chiral flow is maximal, i.e. the onset of streak regression $t = 0$. For calculating the forces driving the large-scale symmetric flow towards the streak, we use $l_h \to \infty$, $\alpha = 1$ and $\eta = 1\,\text{mN}\,\text{h/m}$. As expected from the previous section we find that active forces are mostly localized to the streak and the node (Fig. 2a,b). At the streak, forces mostly point towards the center-line of the streak. At the node, forces are smaller in magnitude than at the streak and mostly point towards the center of the node. Taken together, the forces $\mathbf{f}_{\text{act}}$ calculated from the symmetric flow are consistent with an active contraction of the node and streak tissue, with the streak contracting primarily along the left-right axis. This is consistent with models for streak formation that were recently proposed in [52,53,44].

Next, we calculate the forces underlying chiral flow. As discussed in the previous section, the measured flow field suggests a finite hydrodynamic length $l_h \sim 50 - 100\mu\text{m}$. We use $l_h = 75\mu\text{m}$, $\alpha = 1$ and $\eta = 1\,\text{mN}\,\text{h/m}$. As discussed in the main text, we find that the forces calculated from the measured chiral flow are localized to the node tissue. From this, we calculate the net force $\mathbf{F}_{\text{node}}$ and the net torque $\Gamma_{\text{node}}$ that the node generates, i.e.

$$\mathbf{F}_{\text{node}} = \int_{|\mathbf{r}|\leq 100\mu\text{m}} \mathrm{d}S\, \mathbf{f}_{\text{act}}, \quad \Gamma_{\text{node}} = \int_{|\mathbf{r}|\leq 100\mu\text{m}} \mathrm{d}S\, \mathbf{z}\cdot(\mathbf{r}\times\mathbf{f}_{\text{act}})\,, \tag{S38}$$

where $\mathbf{r}$ is the position vector relative to the center of the node. To this end, we calculate the average value of the integrand using the calculated $\mathbf{f}_{\text{act}}$ for grid points with $|\mathbf{r}| \leq 100\mu\text{m}$ and multiply this mean value with the area of the circle. We obtain $\mathbf{F}_{\text{node}} = 0.05\mu\text{N}$ and $\Gamma_{\text{node}} = 5.8\mu\text{N}\mu\text{m}$. The net force and torque account for most of the force field. This suggests that a net force and torque generated by the node underlies chiral flow. In section 2.5, we discuss a model where chiral flows are driven by an active torque density and the non-vanishing $\mathbf{F}_{\text{node}}$ we obtain here is a consequence of the rigidity of the streak.

One may wonder, however, how confident we can be about $\mathbf{F}_{\text{node}}$ and $\Gamma_{\text{node}}$ given that we do not know the material properties of the node and the streak. Using Eq. S19, we can distinguish between a viscous and a friction contribution to the net force and torque. We find that for the torque, the viscous contribution dominates over the friction contribution ($3.7\mu$N$\mu$m vs $2.1\mu$N$\mu$m for $l_h = 75\mu$m). Intriguingly, we can rewrite the viscous contribution using Gauss theorem:

$$F_i^{\text{node}} = -\int_{|\mathbf{r}|=100\mu\text{m}} \mathrm{d}l\, t_{ji}^{\text{visc}} \hat{r}_j + \int_{|\mathbf{r}|\leq 100\mu\text{m}} \mathrm{d}S\, \gamma v_i, \tag{S39}$$

$$\Gamma_{\text{node}} = -\int_{|\mathbf{r}|=100\mu\text{m}} \mathrm{d}l\, r t_{ji}^{\text{visc}} \hat{\theta}_j + \int_{|\mathbf{r}|\leq 100\mu\text{m}} \mathrm{d}S\, \gamma r(\hat{\theta}\cdot\mathbf{v}), \tag{S40}$$

where $\hat{\mathbf{r}}$ and $\hat{\boldsymbol{\theta}}$ are the radial and azimuthal unit vectors, respectively. Here, the viscous term is a boundary integral. Importantly, this is valid also for gradients of viscosity inside the enclosed area of the node. Thus, the force and torque resulting from viscous forces is independent of the material parameters within the node, even though we calculate from the experimental data using a surface integral. The viscous force and torque does depend, however, on the viscosity of the streak. In general, an increased viscosity of the streak tissue could imply that the net force and torque, which the node generates, vanishes. In such a scenario, the streak would provide the torque and force the node exerts onto the epiblast. However, the laser-ablation experiments imply that the node drives its own rotation, implying a net torque generated at the node (Fig. 2). Also observations of streak ruptures indicate that mechanical integrity of the streak is not necessary for the rotation of the node (data not shown). Thus, we conclude that the node generates a torque to drive a chiral flow of the surrounding epiblast tissue.

This is striking, since such a torque requires mechanical coupling to a substrate. To see this explicitly, we evaluate Eq. S20, considering a model where all sources of mechanical activity vanish everywhere except within the node tissue. Thereby we find

$$\Gamma_{\text{node}} = \int_{\mathbf{r}\leq 100\mu\text{m}} \mathrm{d}S\, \left[\tau_{\text{act}} + \mathbf{z}\cdot(\mathbf{r}\times\mathbf{f}_{\text{act}})\right]. \tag{S41}$$

Thus, a non-vanishing torque $\Gamma_{\text{node}}$ implies a non-vanishing torque or force density generated with respect to the underlying substrate.

#### 2.4 Torque balance of the node upon circular laser cuts

Angular momentum implies that all the torques and forces exerted on a piece of tissue have to sum up to zero, as inertia effects are negligible. In the following we discuss the balance of torques at the node before and after circular laser cuts that mechanically isolate the Hensen's node from the surrounding epiblast (see Fig. 2e-h). We consider two scenarios. In Scenario I chiral flow is driven by an active torque $\Gamma_{\text{act}}$ that an underlying substrate exerts on the dorsal tissue of the node. In Scenario II, chiral flow results from the node propelling itself relative to the surrounding epiblast, while mechanical coupling to the substrate slows down chiral flow. The analysis of the spatial profile of chiral flow suggests Scenario I as discussed in the previous section.

In Scenario I, the substrate exerts a torque $\Gamma_{\mathrm{act}} + \Gamma_{\mathrm{fric,node}} > 0$ on the dorsal node tissue, where $\Gamma_{\mathrm{act}} = \int \mathrm{d}S\tau_{\mathrm{act}}$ and $\Gamma_{\mathrm{fric,node}} \sim -\gamma\Omega_{\mathrm{node,rel}}R^2_{\mathrm{node}}$. This net torque is balanced by mechanical interaction with the surrounding epiblast, i.e. the node exerts the torque $\Gamma_{\mathrm{in-plane}} = \Gamma_{\mathrm{act}} + \Gamma_{\mathrm{fric,node}}$ on the surrounding epiblast, which drives chiral flow there. This torque is then further transmitted through the epiblast and ultimately to the substrate through friction, i.e. $\Gamma_{\mathrm{fric,epi}} = \Gamma_{\mathrm{in-plane}}$ such that the whole system consisting of node, epiblast and substrate is torque-free. The circular laser cut mechanically isolates the epiblast from the node such that $\Gamma_{\mathrm{in-plane}} = 0$. Leaving everything unchanged, the torque that the substrate exerts on the dorsal node tissue would no longer be balanced as $\Gamma_{\mathrm{act}} + \Gamma_{\mathrm{fric,node}} > 0$, yielding an accelerated rotation of the node. On the viscous time scale, i.e. hours, this implies that the node now rotates with an increased velocity $\Omega'_{\mathrm{node,rel}}$ such that $\Gamma_{\mathrm{act}} + \Gamma'_{\mathrm{fric,node}} = 0$. On the time scale of seconds, the entire system, i.e. the epiblast, the underlying tissue and the mechanical links between tissue layers, behaves as an elastic material (see the above discussion on the viscosity of the epiblast). In this regime, an imbalance of torques yields an immediate deformation of the tissue such that elastic stresses amount to a torque dipole between dorsal node tissue and underlying substrate, balancing the active torque dipole ($\Gamma_{\mathrm{act}}$). Such a deformation yields an increase in elastic energy, which is provided by the work the active torque dipole performs on the tissue. As this work has to be greater than zero, the deformation has to amount to a rotation of the epiblast in the direction of chiral flow relative to the underlying substrate, as observed in the experiment (Fig. 2g). In the surrounding epiblast, in contrast, mechanical isolation from the node yields a relaxation of elastic stresses such that $\Gamma_{\mathrm{in-plane}} = 0$. Such a relaxation, yielding a release of elastic energy, amounts to a rotation of the surrounding epiblast opposite to the direction of chiral flow, as observed in the experiment (Fig. 2h).

In Scenario II, the substrate exerts a torque $\Gamma_{\mathrm{fric,node}} < 0$ on the dorsal node tissue, which is balanced by torque $-\Gamma_{\mathrm{in-plane}} > 0$ that the surrounding epiblast exerts on the node. Thus, the roles of node and epiblast are reversed compared with Scenario I. Thus, Scenario II implies that upon a circular laser cut the surrounding epiblast rotates in the direction of chiral flow, whereas the node rotates in the opposite direction, contradicting experimental observations (Fig. 2f-h).

##### 2.5 A minimal model of chiral flow

In the following, we propose a minimal active fluid model that quantitatively captures the chiral flow in the avian epiblast. We consider a fluid film coupled to a substrate as introduced in section 2.1. To capture chiral flows, we allow for an active torque density $\tau_{\mathrm{act}}$, that is non-vanishing only up to a distance of $125\mu$m from the node center. We consider the scalar field $\tau_{\mathrm{act}}$ to be left-right symmetric but allow for an asymmetry along the AP axis, since the tissue structure, as well as the concentration of morphogens, are clearly asymmetric along this axis [11]. A simple smooth function that fulfills all these requirements is a linear combination of the first two eigenmodes of the Laplace operator for Dirichlet boundary conditions on a circle:

$$\tau_{\text{act}} = \tau_0 J_0(r/\lambda_0) + \tau_1 J_1(r/\lambda_1)\cos(\theta), \tag{S42}$$

where $J_m$ are the Bessel functions of the first kind and $r, \theta$ are polar coordinates with respect to the node center with $\theta = 0$ coinciding with the mid-line anterior to the node. $\lambda_0 \approx 41.6\mu$m and $\lambda_1 \approx 26.1\mu$m are length scales defined by the boundary condition $\tau_{\text{act}}(r = 100\mu\text{m}) = 0$. $\tau_0$ and $\tau_1$ are coefficients that we fit to the measured flow field as explained in the following.

For simplicity, we use $\alpha = 1$ and $l_h = 75\mu$m throughout the plane of the tissue. Of course, the tissue architecture of the streak is markedly different from the surrounding epiblast. In particular, cell debris and extracellular matrix at the center of the streak prevents cells from crossing the mid-line [86]. Also the flow field we measure implies that ingression is limited to a band on the left and the right side of the streak center (Fig. S8). To capture this mid-line barrier we consider a 10-fold increase in tissue viscosity at the streak, specifically in a rectangle of width $100\mu$m centered at the streak center-line, which covers the streak up to distance of $50\mu$m from the node center (Fig. 2i). As we keep $l_h$ constant, this translates also into in a 10-fold increase in the friction coefficient $\gamma$. We interpret this as $\eta$, $\alpha\eta$ and $\gamma$ all having the same origin, namely the 3D visco-elasticity of the streak tissue.

The flow field is governed by the force balance equation S19 with

$$f_i^{\text{act}} = \epsilon_{ij}\partial_j \tau_{\text{act}}. \tag{S43}$$

We solve this differential equation numerically on a circular domain with radius $400\mu$m. To this end, we calculate derivatives using finite differences on a staggered square grid of size $100 \times 100$ with grid-spacing $8.4\mu$m and the grids for velocity staggered with respect to the grid for torque density and stresses by $4.2\mu$m (see also [77]). With this we obtain the components of the flow field generated by $\tau_0$ and $\tau_1$. We then determine $\tau_0$ and $\tau_1$ by linear regression. To this end we interpolate the measured flow field at the grid points of the numerical solutions. We then determine $\tau_0$ and $\tau_1$ such that the mean squared residual of the measured chiral flow is minimal for data points in a distance of up to $200\mu$m of the node center, excluding data points within a distance of $25\mu$m to the streak center, where the measured flow field tends to be less reliable due to debris. Thereby, we obtain

$$\tau_0/\eta \approx 0.575 h^{-1}, \quad \tau_1/\eta = -0.318 h^{-1}, \tag{S44}$$

which we use in Fig. 2 in the main text. Here $\eta = 1\,\text{mN}\,\text{h/m}$ is the viscosity of the epiblast.

#### 2.6 Mechanical model of tissue layer coupling

As discussed in section 2.3, we inferred a hydrodynamic length $l_h = \sqrt{\eta/\gamma}$ by analyzing the measured flow field $\mathbf{v}$ in the dorsal epiblast, using a model of a homogeneous fluid film coupled to a rigid substrate (Eq. S21). We analyzed the L/R symmetric component $\mathbf{v}_{\mathrm{sym}}$ (Eq. S5, Fig. S6) and the L/R antisymmetric component $\mathbf{v}_{\mathrm{antisym}}$ (Eq. S6, Fig. S7), separately. Strikingly, we found a small hydrodynamic length $l_h = 50 - 100\mu\mathrm{m}$ when analyzing $\mathbf{v}_{\mathrm{antisym}}$ corresponding to the chiral flow at the node. The large-scale flows towards the streak $\mathbf{v}_{\mathrm{sym}}$, in contrast, are associated with a large hydrodynamic length $l_h > 300\mu\mathrm{m}$ on the order of the system size, consistent with earlier studies [52, 44,53]. These distinct hydrodynamic lengths imply that the mechanical nature of chiral flow is distinct from the large-scale symmetric flows. This is puzzling, since both flows happen at the same time in the same tissue, only our analysis separates these flow components. As discussed in the main text, perturbation experiments reveal that the mechanical coupling between the epiblast and the underlying meso/endoderm facilitates chiral flows but not large-scale tissue movements. In the following, we discuss how mechanical coupling between the tissue layers can yield distinct hydrodynamic lengths for different flow components.

We describe the dorsal epiblast and the underlying ventral tissue as two fluid films with distinct viscosities $\eta_d$ and $\eta_v$, respectively . The two fluid films are coupled in terms of a force density

$$\mathbf{f}_{v \to d} = -\gamma_{dv}(\mathbf{v} - \mathbf{v}_v) \tag{S45}$$

that the ventral tissue exerts onto the dorsal tissue. Here, $\mathbf{v}$ and $\mathbf{v}_v$ are the flow fields in the dorsal and ventral tissue, respectively. In the experiment only $\mathbf{v}$ is measured. $\mathbf{f}_{v \to d}$ is balanced by a force density $\mathbf{f}_{d \to v}$ that the dorsal tissue exerts on the ventral tissue. Furthermore, the fluid films are coupled by an active torque density $\tau_{dv}$ that rotates the dorsal relative to the ventral tissue. To capture the flux towards the streak we consider an active symmetric stress $t_{ij}^{\mathrm{act,d}}$ in the dorsal tissue only. As before, we consider gradients of mechanical activity (i.e. $\tau_{dv}$ and $t_{ij}^{\mathrm{act,d}}$) to vanish outside the streak and the node region. For simplicity, we consider the viscosities $\eta_d, \eta_v$ to be constant throughout the plane and consider a single value of $\alpha$ for the dorsal and ventral tissue. With this, tangential force balance (Eq. S19) yields the following two coupled differential equations for the flow fields:

$$\eta_d \Delta \mathbf{v} + \alpha \eta_d \boldsymbol{\nabla}(\operatorname{div} \mathbf{v}) = \gamma_{dv}(\mathbf{v} - \mathbf{v}_v) - \partial_j \mathbf{t}_j^{\mathrm{act,d}} - \mathbf{z} \times \boldsymbol{\nabla} \tau_{dv} \tag{S46}$$

$$\eta_v \Delta \mathbf{v}_v + \alpha \eta_v \boldsymbol{\nabla}(\operatorname{div} \mathbf{v}_v) = -\gamma_{dv}(\mathbf{v} - \mathbf{v}_v) + \mathbf{z} \times \boldsymbol{\nabla} \tau_{dv} \tag{S47}$$

By adding and subtracting the above equations divided by the viscosities, we rewrite them as equations of the total tissue flow $\mathbf{v}+\mathbf{v}_v$ and the relative flow $\mathbf{v}-\mathbf{v}_v$:

$$\Delta(\mathbf{v}+\mathbf{v}_v)+\alpha\boldsymbol{\nabla}\left[\operatorname{div}(\mathbf{v}+\mathbf{v}_v)\right]=\frac{\gamma_{dv}}{\eta_d}\frac{\eta_v-\eta_d}{\eta_v}(\mathbf{v}-\mathbf{v}_v)-\frac{1}{\eta_d}\partial_j\mathbf{t}_j^{\mathrm{act,d}}-2\frac{\eta_v-\eta_d}{\eta_d\eta_v}(\mathbf{z}\times\boldsymbol{\nabla}\tau_{dv}) \quad \text{(S48)}$$

$$\Delta(\mathbf{v}-\mathbf{v}_v)+\alpha\boldsymbol{\nabla}\left[\operatorname{div}(\mathbf{v}-\mathbf{v}_v)\right]=\frac{\gamma_{dv}}{\eta_d}\frac{\eta_v+\eta_d}{\eta_v}(\mathbf{v}-\mathbf{v}_v)-\frac{1}{\eta_d}\partial_j\mathbf{t}_j^{\mathrm{act,d}}-2\frac{\eta_d+\eta_v}{\eta_d\eta_v}(\mathbf{z}\times\boldsymbol{\nabla}\tau_{dv}). \quad \text{(S49)}$$

We observe that the differential equation for the $\mathbf{v}-\mathbf{v}_v$ (Eq. S49) is in general independent of $\mathbf{v}+\mathbf{v}_v$. Furthermore, Eq. S49 defines a hydrodynamic length

$$l_{h,dv}=\sqrt{\frac{\eta_d\eta_v}{\gamma_{dv}(\eta_v+\eta_d)}}. \quad \text{(S50)}$$

Outside the streak and the node, gradients of mechanical activity (i.e. $\tau_{dv}$ and $\mathbf{t}_j^{\mathrm{act,d}}$) vanish and thus Eq. S49 yields an exponential decay of the relative flow $\mathbf{v}-\mathbf{v}_v$, i.e.

$$|\mathbf{v}-\mathbf{v}_v|\sim e^{-r_{\mathrm{streak}}/l_{h,dv}}, \quad \text{(S51)}$$

where $r_{\mathrm{streak}}$ is the distance to the streak center. Farther away from node and streak ($r_{\mathrm{streak}}\gg l_{h,dv}$), $\mathbf{v}-\mathbf{v}_v$ vanishes and ventral and dorsal tissue layers move together as one fluid film with infinite hydrodynamic length, as Eq. S48 becomes equivalent to Eq. S21 with $l_h\to\infty$.

It is instructive to consider the scenario of fluid films with equal viscosity ($\eta_d=\eta_v$). Then, the equations for $(\mathbf{v}+\mathbf{v}_v)$ and $(\mathbf{v}-\mathbf{v}_v)$ decouple everywhere (see Eq. S48, S49 with $\eta_v-\eta_d=0$). In such a system, only the active symmetric stress $t_{ij}^{\mathrm{act,d}}$ drives a total flow $\mathbf{v}+\mathbf{v}_v$. Furthermore, $\mathbf{v}+\mathbf{v}_v$ is the flow of fluid with infinite hydrodynamic length, implying that $\mathbf{v}+\mathbf{v}_v$ spans the entire embryo. The active torque $\tau_{dv}$, in contrast, drives only a relative movement $(\mathbf{v}-\mathbf{v}_v)$ of the tissue layers, which decays on the hydrodynamic length scale $l_{h,dv}$ away from the node and the streak.

In this picture, gradients of $t_{ij}^{\mathrm{act,d}}$ at the streak drive the observed large-scale L/R-symmetric flow $\mathbf{v}_{\mathrm{sym}}$. While the L/R symmetric component $\mathbf{v}_{\mathrm{sym}}$ of $\mathbf{v}$ generally contains contains contributions from both $(\mathbf{v}+\mathbf{v}_v)$ and $(\mathbf{v}-\mathbf{v}_v)$, $(\mathbf{v}+\mathbf{v}_v)$ dominates away from the streak due to the sharp decay of $\mathbf{v}-\mathbf{v}_v$ (Eq. S51). This explains the large hydrodynamic length we infer from the measured spatial profile of $\mathbf{v}_{\mathrm{sym}}$. In contrast, we explain the chiral flow $\mathbf{v}_{\mathrm{antisym}}$ as a result of an active torque density $\tau_{\mathrm{dv}}$ that vanishes away from the node (see previous section). Thus, the measured L/R-antisymmetric component $\mathbf{v}_{\mathrm{antisym}}$ of $\mathbf{v}$ corresponds solely to a relative movement $(\mathbf{v}-\mathbf{v}_v)$ of the tissue layers. This explains the observed exponential decay of $\mathbf{v}_{\mathrm{antisym}}$ away from the node (Fig. S7). Thus, we interpret the small hydrodynamic length $l_h$ inferred from $\mathbf{v}_{\mathrm{antisym}}$ as the hydrodynamic length $l_{h,dv}$ that results from the mechanical coupling of tissue layers, i.e. $l_{h,dv}=l_h=50-100\mu$m. We conclude that a finite hydrodynamic length of chiral flow arises naturally, when chiral flow is driven by a torque dipole between mechanically coupled layers. As discussed in the main text, perturbation experiments show that mechanical coupling of tissue layers does indeed facilitate the generation of chiral flow (Fig. 3).

Notably, our model provides two predictions: First, chiral flow amounts to a relative movement of tissue layers. This asks for future studies to investigate the 3D profile of chiral flow across tissue layers. Second, dorsal and ventral tissue layers move together as one fluid on the embryonic scale. Strikingly such a common movement of epiblast and underlying tissue has indeed recently been observed, though at an earlier stage of development [57].

While our model is instructive, we do not expect it to capture the full complexity of the embryo. In particular, we expect that ventral and dorsal tissue are mechanical distinct (i.e. $\eta_d \neq \eta_v$). We note, however, that also in such a more complex scenario the active torque density $\tau_{dv}$ driving chiral flow contributes primarily to the relative movement $(\mathbf{v} - \mathbf{v}_v)$ in contrast to the active stress $t_{ij}^{\mathrm{act,d}}$. This can be seen from Eq. S48,S49 using that for any $\eta_v, \eta_d > 0$, $|\eta_v - \eta_d| < |\eta_v + \eta_d|$. Thus, the qualitative arguments discussed above remain valid. Furthermore, we expect the streak tissue to be mechanically distinct. In the previous section, we already discussed the evidence for an increased viscosity $(\eta_d, \eta_v)$ at the streak. On top of that, we also expect gradients of $\gamma_{dv}$. In contrast to the stages considered in [57], during the later stages of development we study here, the invading mesoderm decouples the epiblast from the hypoblast and future endoderm. Thus, mechanical coupling of tissue layers (i.e. $\gamma_{dv}$) may become negligible away from the streak. Note in particular that local degradation of basal membrane supports the idea of mechanical coupling occurring only at the node and streak region (see Fig. S10). This may allow for large-scale movements $\mathbf{v} - \mathbf{v}_v$ of the dorsal relative to the ventral layer. Importantly, active torques $\tau_{dv}$ can only arise where the layers are tightly coupled. Thus, chiral flow can only arise where $\gamma_{dv}$ is non-negligible and thus $l_{h,dv}$ is small. In such a scenario, the apparent hydrodynamic length of chiral flow, being limited to the node area, would still be distinct from the large-scale left-right symmetric flow. Thus, also in such a more complex scenario, the distinct hydrodynamic length we infer from chiral flow is a consequence of the distinct mechanical nature of chiral flow $(\mathbf{v}_{\mathrm{antisym}})$ as compared to the large-scale flow $\mathbf{v}_{\mathrm{sym}}$, also called polonaise flow: Chiral flow arises from a torque dipole between tissue layers, whereas large-scale polonaise flow arise from force-dipoles within the dorsal tissue layer.

#### 2.7 Streak kinking as a consequence of chiral flow

While the primitive streak appears as a straight left-right symmetric line up to the point of streak regression (see Fig. 1d,k), the anterior tip of the streak kinks towards the left at latter time points (Fig. 1g,k). We hypothesize that this kinking is an immediate consequence of chiral flow due to advection. As shown in Fig. 2i-m, we can account for chiral flow by considering a fluid model where flows are driven by active torque dipoles at the node. We asked, if this model can also account for the kinking of this streak. To this end, we consider an initially straight line along the mid-line ($x = 0$) consisting of 400 points between $y = -400\mu$m and $y = +400\mu$m with respect to the node center. We then advect each point using the chiral flow calculated from the model (Fig. 2j) over a time of 6h using simple Euler integration. At each integration time point we interpolate the static flow field (Fig. 2j) at the current position of the point. We observe that the mid-line immediately posterior to the node center kinks leftward similarly the kinking we observe for the anterior tip of the streak (Fig. S9). We quantify the kinking angle of the advected line by calculating the angle with the mid-line of the line connecting the points that are initially immediately anterior and posterior to the node center, yielding $\theta_{\text{kink}} = 44.6°$. Note that this should be interpreted only as a rough estimate, since we do not take into account here that a) the left-right-symmetric flow contributes to the advection once the line is kinked and b) chiral flow changes over time likely also due to advection of torque generators (Fig. S4). However, these effects amount to higher order corrections, such that we expect that the qualitative result of our calculation, the leftward kinking, remains valid.

#### 2.8 On mechanical coupling to the vitelline membrane

The laser cut experiments provide evidence that chiral flow in the epiblast is driven by a torque generated at the node due to mechanical coupling to a substrate. In principle, this substrate could lie on the dorsal or ventral side of the epiblast. The epiblast is situated between the vitelline membrane on the dorsal side and the meso/endoderm on the ventral side (Fig.1b). Hence either side could act as the substrate that sustains the counter-torque. However, it has been previously shown that the epiblast is only mechanically attached to the vitelline membrane at the outermost edge of the growing extra-embryonic tissue [87,88,89,90]. We would therefore not expect a strong mechanical interaction between the epiblast and the vitelline membrane in the area of the Hensen's node. To test if mechanical coupling to the vitelline membrane is required for chiral flow, we inserted a small hole in the vitelline membrane directly above the Hensen's node. This manipulation did not perturb chiral flow or streak kinking (Fig.3f-i and S5). Thus, mechanical coupling of the epiblast to the vitelline membrane on the dorsal (or equivalently apical) side is not required for chiral flow at the node.

However, this finding does not contradict that in general the vitelline membrane is a compatible substrate, with which cells can interact tightly. While we find that mechanical coupling to the ventral tissue is required for chiral flow, we also find that it can be replaced by a vitelline membrane without perturbing chiral flow (Fig. 3). In this condition (ventral tissue replaced), the inserted vitelline membrane is in contact with the basal (=ventral) side of the epiblast, in contrast to the unperturbed embryo. Thus, we conclude that chiral flow relies on mechanical coupling to a basal or ventral substrate.

### 3 Supplementary Figures

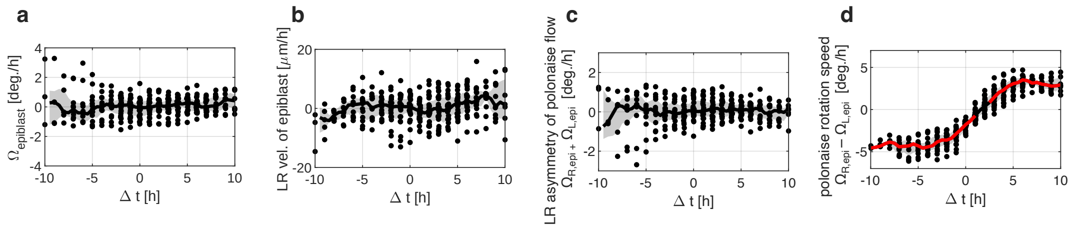

**Figure S1: Quantification of left-right asymmetries of large-scale flow.** We quantify the rotation speed (**a**, Eq. S3) and leftward velocity (**b**, Eq. S3) of the epiblast surrounding the node in a distance between 350$\mu$m and 600$\mu$m in the lab frame, which shows no consistent chiral net rotation or translation of the epiblast. Furthermore we quantify polonaise flow (**d**) and its left-right asymmetry (**c**) in terms of an average vorticity (Eq. S9), where we again do not observe a consistent left-right asymmetry.

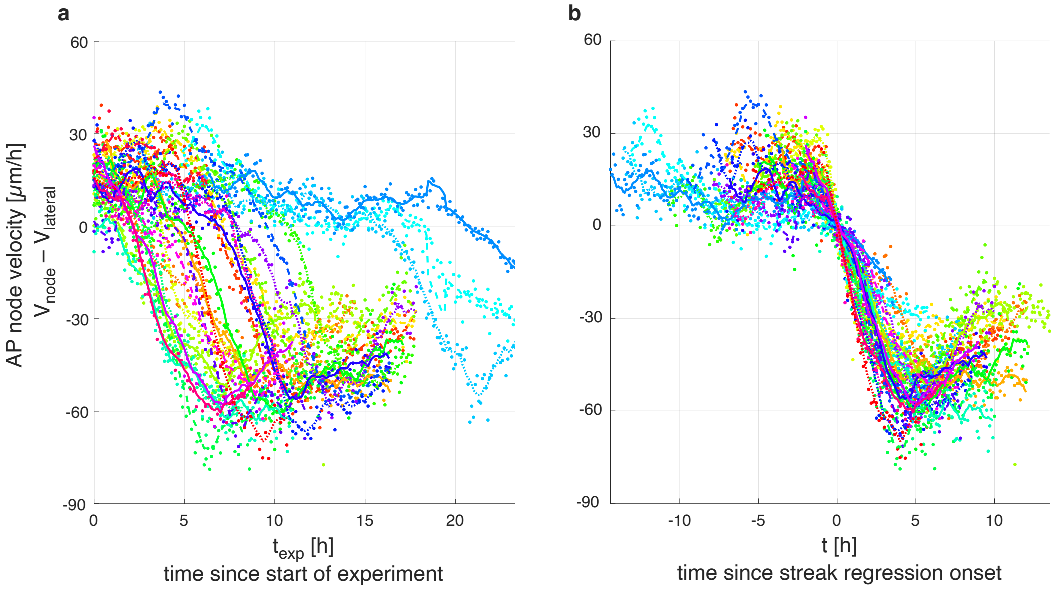

**Figure S2: Aligning embryos in time.** **a**: anterior-ward velocity of the node relative to lateral epiblast (see section 1.2.3) as a function of time. Different colors denote different embryos (n=25, all under control conditions) with dots denoting single time points and the lines denoting the moving average using a 1h time window. From the moving average we determine the time point of streak regression onset (t=0h) as described in section 1.2.3. In **b**, the anterior-ward velocity of the node is plotted as in **a** but as a function of the time $t$ since streak regression onset.

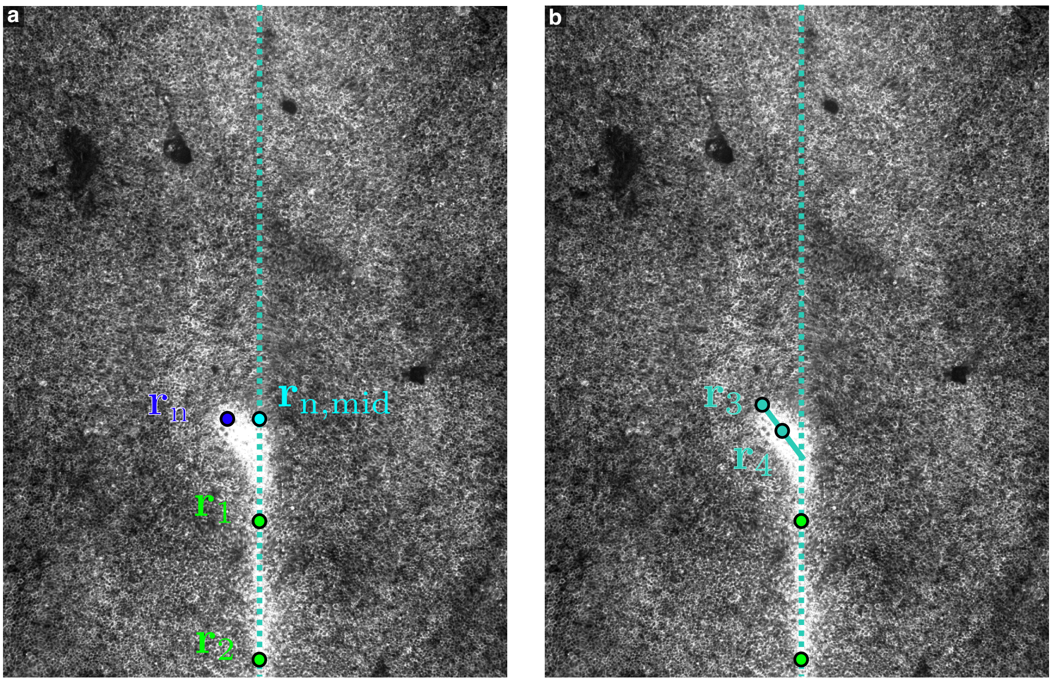

**Figure S3: Manual annotation of streak and node.** **a** illustrates the annotation of the streak orientation (in terms of $\mathbf{r}_1$ and $\mathbf{r}_2$) and node position ($\mathbf{r}_\mathrm{n}$) as explained in section 1.2.3. **b** illustrates the annotation of the kink angle, where we annotate the long axis of the node in terms of $\mathbf{r}_3$ and $\mathbf{r}_4$ as explained in section 1.2.7.

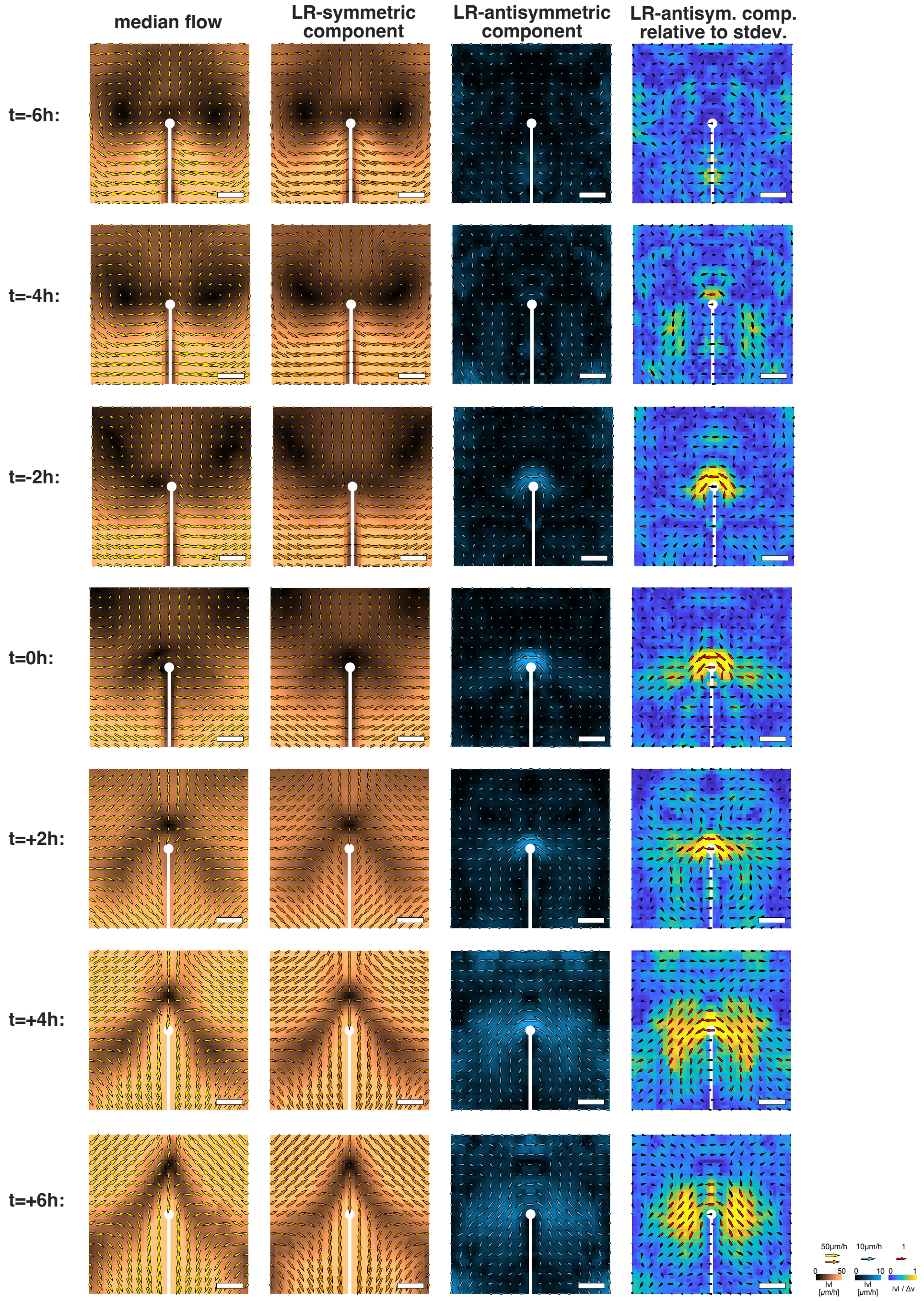

**Figure S4: Median flow field as a function of time.** Median was calculated across n=12,14,22,25,25,23,19 unperturbed embryos at time points t=-6,-4,-2,0+2,+4 hours with respect to streak regression onset, respectively. Columns 1, 2 and 3 from the left show median total flow, median left-right symmetric flow and median left-right antisymmetric flow, respectively, where arrows denote length and magnitude of tissue velocity and the color additionally illustrates the magnitude. In the right column, we plot the median left-right antisymmetric flow normalized with respect to the experimental standard deviation of the tissue velocity, but otherwise plotted analogous to the other columns and Fig. 1l-n. Red vectors denote data points where at least one of the components ($v_x$ or $v_y$) differs significantly from zero at a significance value of p=0.01 using a Wilcoxon rank sum test. Scale bars are 200μm.

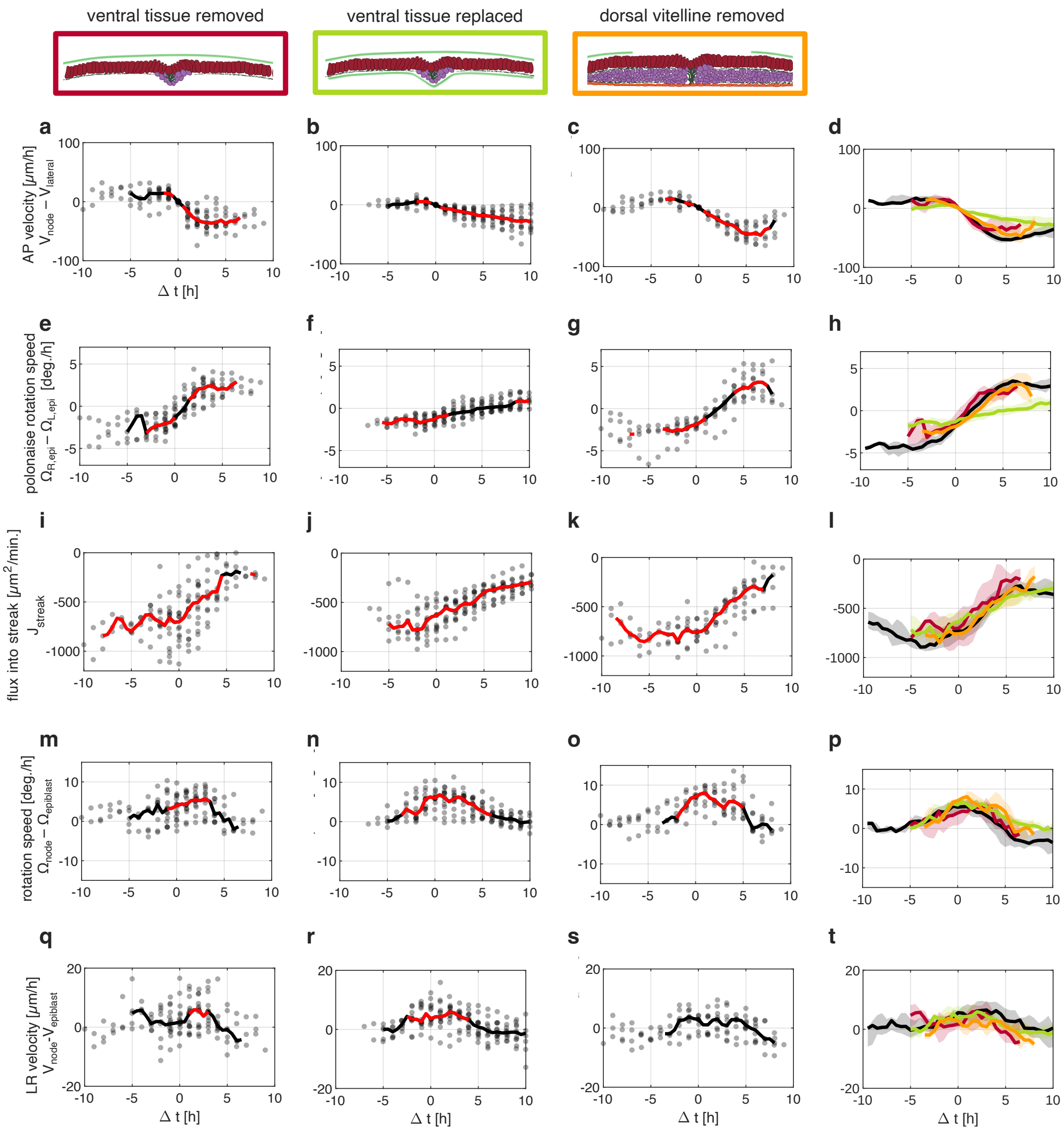

**Figure S5: Quantification of tissue flow in perturbed embryos.** First 3 columns (**a-c,e-g,i-k,m-o,q-s**): As in Fig. 1h-j, gray dots are single embryos, solid line denotes the median with red color denoting time points where the median is significantly different from 0 ($p < 0.01$, Wilcoxon signed rank test). $t$ denotes the time with respect to streak regression onset at $t = 0$. Schematics denote the experimental condition of the data in the column underneath. Right column **d,h,l,p,t**: solid lines denote embryo median, shaded areas denote the range of the central 50 percent of the data. Colors denote experimental condition as in Fig. 3g-i. **a-d**: AP velocity of node with respect to lateral epiblast as defined in section 1.2.3 and plotted in Fig. 1h. **e-h**: symmetric vorticity of epiblast $\Omega_L + \Omega_R$ (Eq. S9). **i-l**: flux into streak (Eq. S10). **m-p** Rotation of node relative to surrounding epiblast tissue (Eq. S7). **q-t**: leftward velocity of node relative to surrounding tissue (Eq. S8). Taken together, we observe that removal of ventral tissue delays and inhibits chiral flow (**m,q**), but leaves the non-chiral flow largely unaffected (**a,e,i**). In contrast, replacement of the ventral tissue by a surrogate substrate (vitelline membrane) slows down large scale flows (**b,f**), whereas timing and magnitude of chiral flow are comparable to unperturbed embryos **n,r**.

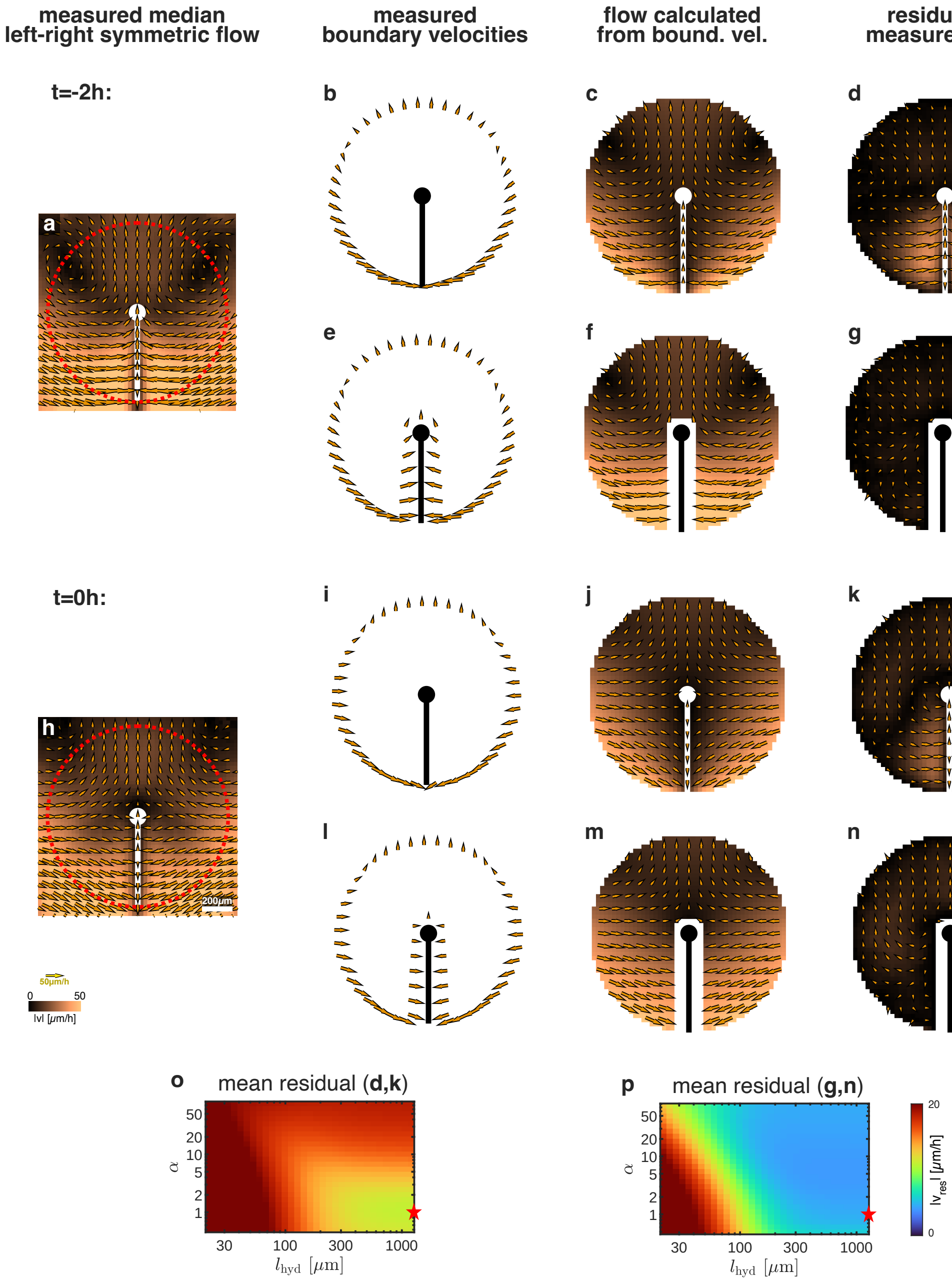

**Figure S6: Boundary analysis of left-right symmetric flow. a**: median left-right symmetric component of the measured flow field at $t = -2h$ with respect to streak regression onset. Red circle denotes an outer boundary at a distance $r = 600\mu$m with respect to the node (white circle). **b** flow field (same as in **a**) evaluated at this circle. **c**: flow field calculated from the velocity boundary condition in **b** using a fluid model with $\alpha = 1$,$l_h = 1200\mu$m. **d**: residual of the flow field in **a** after subtracting the calculated flow field in **c**. **e** flow field (same as in **a**) evaluated at an outer boundary as in **b** and at a boundary lining the streak and the node at a distance of $100 - 125\mu m$ with respect to the streak center. **f,g**: analogous to **c,d** but using the boundary velocities in **e**. **h-n**: analogous to **a-g** but using the flow field at $t = 0$ as in Fig. 1l. **o**: root mean squared of the residual as shown in **d,k**, averaged over time points $\{-3h, -2h, -1h, 0h\}$, for a range of material parameters $\alpha, l_h$. This yields a measure to what extent the flow field at streak and epiblast can be captured by our fluid model where flows are only driven at the outer boundary. Red star indicates the material parameters ($\alpha = 1, l_h = 1200\mu$m) used in **c,d,j,k**. **p**: analogous to **o** but for a boundary that excludes the streak and the node as in **e-g,l-n**. This shows that the flow field away from the streak and the node can be captured by a fluid model with $l_h \geq 300\mu$m.

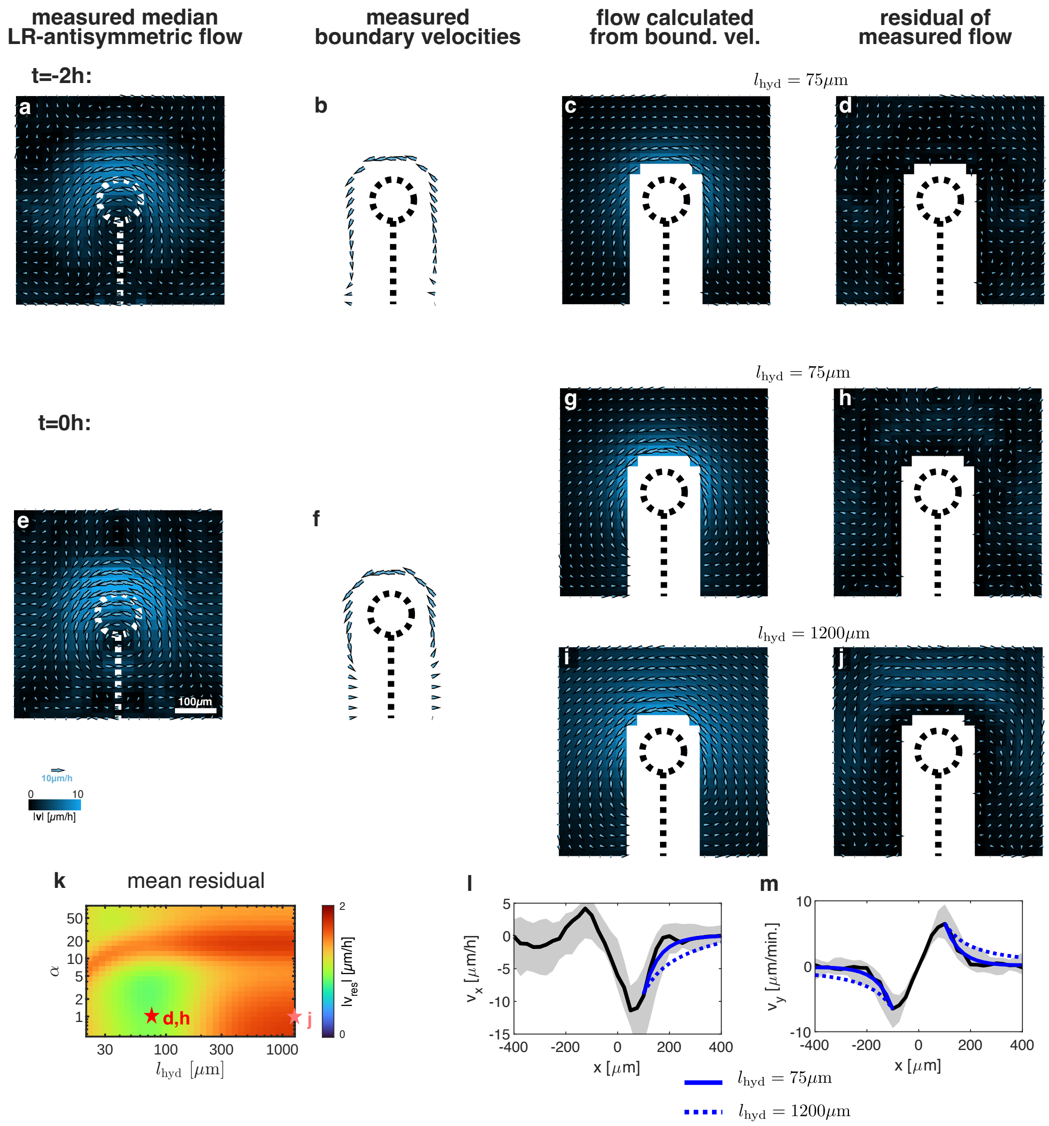

**Figure S7: Boundary analysis of left-right antisymmetric flow.** **a**: median left-right antisymmetric flow field around the node as in Fig. 1n, but at $t = -2h$. **b** flow field in **a** evaluated at a boundary lining the streak and the node at a distance of $100 - 125\mu$m with respect to the node center. **c** flow field calculated from velocity boundary condition consisting of the boundary shown in **b** and an outer circle as in Fig S7a,b where the measured flow field in **a** is evaluated. Material parameters: $\alpha = 1, l_h = 75\mu$m. **d**: residual of the flow field in **a** after subtracting the calculated flow field in **c**. **e-h**: analogous to **a-d** but at $t = 0h$. Note that **e** is identical to Fig. 1n in the main text. **i,j** analogous to **g,h** but using an effectively infinite hydrodynamic length $l_h = 1200\mu$m. **k**: root mean squared of the residual as shown in **d,h,j**, averaged over time points $\{-3h, -2h, -1h, 0h\}$, for a range of material parameters $\alpha, l_h$, taking into account only data points within a distance of $300\mu$m with respect to the node center. This yields a measure to what extent the decay of chiral flow away from the node can be captured by a passive fluid model of the epiblast. Red star indicates the material parameters ($\alpha = 1, l_h = 75\mu$m) used in **c,d,g,h**, as well as in Fig. 2,d in the main text. Pink star indicates the material parameters ($\alpha = 1, l_h = 1200\mu$m) used in **i,j**. **l**:Plot of the AP velocity ($v_y$) at $t = 0$ as function of the left-right position as in Fig. 2l. We compare the calculated flow fields for small hydrodynamic length (blue solid line, as in **g**) and large hydrodynamic length (blue dashed line, as in **i**) with the experimentally measured flow field (black line indicating the embryo median as in **e** with gray area indicating the 25th to 75th percentile). **m:** Analogous to **l** but for the left-right velocity ($v_x$) across a line through the embryo mid-line streak as in Fig. 1m.

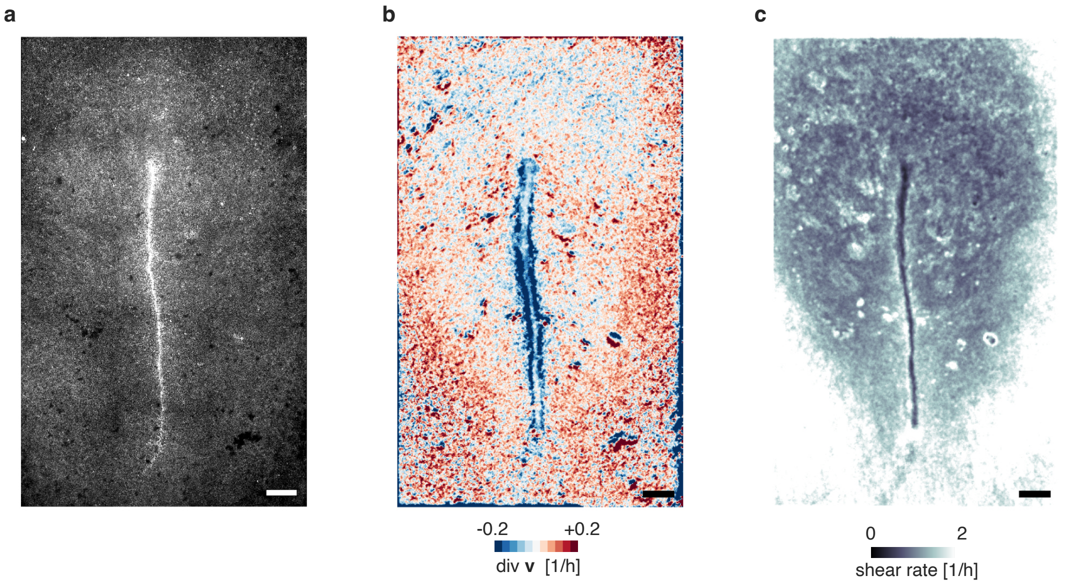

**Figure S8: High resolution spatial profile strain rate at streak.** **a**: microscopy image of epiblast tissue at the time of chiral flow. **b:** divergence of flow field quantified with PIV at a spatial resolution of 5$\mu$m and a time step of 3min. To reduce noise the divergence was averaged over a square of 3x3 data points. Subsequently we obtained the moving median over a time interval of 2h. **c**: analogous to **b** but quantifying the absolute shear rate $\sqrt{(\partial_x v_x - \partial_y v_y)^2 + (\partial_x v_y + \partial_y v_x)^2}$. We observe that the rate of cells leaving the tissue (i.e. negative divergence denoted by blue color in **b**) is maximal in bands to the left and the right of the streak center, while at the streak center the divergence as well as the shear rate (**c**) vanishes. This supports the notion that the streak center is mechanically more rigid than the surrounding epiblast tissue, as assumed in our mechanical model of chiral flow (Fig. 2i). Scale bars: 200$\mu$m

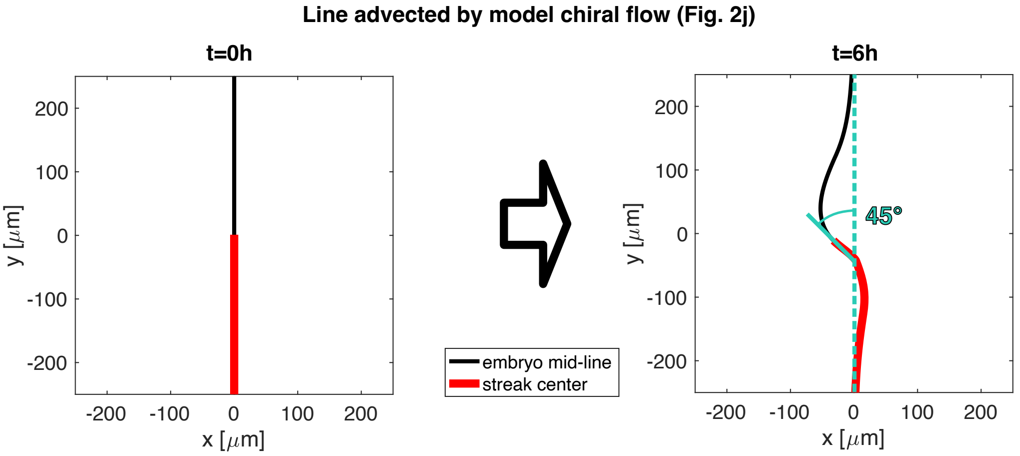

**Figure S9: Modelling streak kinking as a consequence of chiral flow.** We consider an initially straight line advected by the chiral flow in Fig. 2j as described in 2.7. The embryo mid-line (i.e. points initially at $x = 0$) is denoted by a black solid line, the streak (i.e. points initially at $x = 0$, $y \leq 0$ i.e. posterior respect to the node center) is denoted by a red line. Torquoise line illustrate the kinking angle analogous to Fig. 1g.

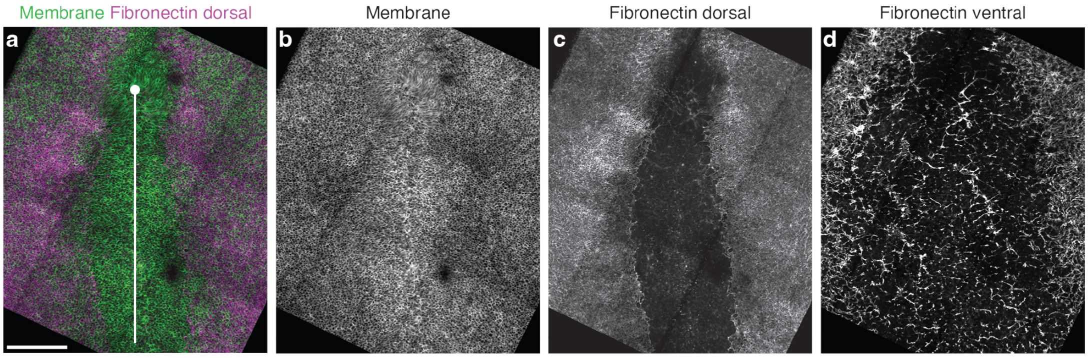

**Figure S10: Fibronectin is absent in the streak region.** Microscopy images of quail epiblast expressing a membrane marker (**a,b**) and stained for Fibronectin (**a,c,d**) - a basal membrane component. Fibronectin is located at the basal side of the epiblast(**c**) where it isolates the epiblast from the mesoderm, and at the basal side of the hypoblast (**d**). Note that in the streak and node region the ECM breaks down and allows for direct interaction between the epiblast and mesoderm cells. White circle and line in **a** marks Hensen's node and primitive streak. Scale bars is 200µm

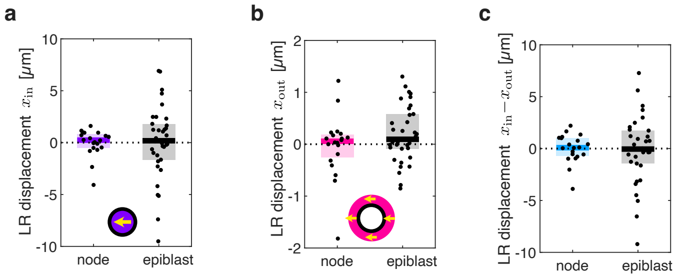

**Figure S11: No net translation of node upon circular laser cut. a:** leftward displacement $x_{\mathrm{in}}$ of tissue enclosed by laser cut (Eq. S13). **b**: leftward displacement $x_{\mathrm{out}}$ of epiblast tissue surrounding the cut region (Eq. S13). **c:** Leftward displacement of cut region relative to surrounding region. As in Fig. 2g,h, displacement was quantified 5s post the beginning of the laser cut. Solid lines indicate median, while shaded area indicates interval from 25th to 75th percentile.

### 4 Supplementary movies

Movie legends:

Mov1: Live imaging of the epiblast of a HH4 quail embryo expressing a GFP-membrane tag capturing the chiral flow around the Hensen's node in the epiblast tissue. T=0 is defined as the onset of streak regression. Scale bar: 200$\mu$m.

Mov2: Live imaging of the epiblast of a HH4 quail embryo expressing a GFP-membrane tag during a circular laser cuts around the Hensen's node. T=0 is the start of the laser cutting procedure. Scale bar:100$\mu$m.

Mov3: Live imaging of the epiblast of a HH4 quail embryo expressing a GFP-membrane tag after the ventral tissue was removed. T=0 is defined as the onset of streak regression. Scale bar: 200$\mu$m.

Mov4: Live imaging of the epiblast of a HH4 quail embryo expressing a GFP-membrane tag after the ventral tissue was replaced by a vitelline membrane.. T=0 is defined as the onset of streak regression. Scale bar: 200$\mu$m.

Mov5: Live imaging of the epiblast of a HH4 quail embryo expressing a GFP-membrane tag after a hole was introduced into the vitelline membrane on top of the Hensen's node. T=0 is defined as the onset of streak regression. Scale bar: 200$\mu$m.
